# United we stand: Plants that physically touch each other are more resilient to excess light stress

**DOI:** 10.1101/2025.09.02.673745

**Authors:** María Ángeles Peláez-Vico, Yosef Fichman, Abdul Ghani, Sameep Dhakal, Ahmad Bereimipour, Rajeev Azad, Stanislaw M. Karpiński, Trupti Joshi, Ron Mittler

## Abstract

Plants use sophisticated signaling networks to communicate with each other. While this process is thought to support the overall health and resilience of plant communities, it could also reflect eavesdropping between plants used for competition. Here we reveal that plants that physically touch each other aboveground are more resilient to excess light stress, and that this phenomenon is dependent on the ability of plants to exchange aboveground electric and H_2_O_2_ signals with each other. Using a mutant that is unable to transfer Ca^2+^/reactive oxygen species (ROS) signals but can transfer electric signals (*hpca1*), as a mediator/connector between different plants, we further separate electric from Ca^2+^/ROS plant-to-plant signals and transcriptional landscapes, and show that the transfer of Ca^2+^/ROS signals, as well as the function of several Ca^2+^/ROS-dependent transcripts, is required for excess light stress acclimation. Our study reveals that plants that live together and physically touch each other establish an aboveground community-wide signaling network that enhances their collective resilience to stress.

**Teaser:** Plants that physically touch and exchange aboveground signals with each other are more resilient to excess light stress.

## Introduction

Plants are vital to most ecosystems on Earth. They establish biological networks with other organisms, as well as communicate with each other in multiple ways (*1*–*3*). Some of the well-studied plant-to-plant (P-T-P) signaling pathways include belowground routes involving direct physical touch between roots, interactions between root-associated arbuscular mycorrhiza networks and other microorganisms, and exchange of different compounds/RNA molecules between roots of different plants (*4*–*7*). In addition, aboveground pathways involving volatiles, acoustic sounds, light reflection/filtering, and physical touch between leaves, are also thought to play a role in P-T-P signaling (*8*–*11*). Although communication between plants was proposed to support the overall health and resilience of different plant communities and ecosystems (*8*, *12*–*14*), direct evidence supporting such a role for this process is scarce. Moreover, eavesdropping between plants, used for competition instead of cooperation, was also proposed as a possible role for P-T-P signaling (*15*, *16*). We recently reported that two plants that physically touch each other aboveground via leaf-to-leaf contacts, or through a parasitic plant that connects them, can exchange electric and reactive oxygen species (ROS) signals between them (*9*, *17*). However, whether this P-T-P signaling process plays a key role in enhancing the resilience of a community of plants to stress, and how P-T-P ROS signals are exchanged between different plants, are currently unknown.

Here we reveal that plants that live together and physically touch each other aboveground are more resilient to excess light (EL) stress compared to plants that do not physically touch each other. We further show that aboveground P-T-P signaling in response to EL stress occurs via the exchange of electric and H_2_O_2_ signals between plants. Using different mutants and RNA-Seq analyses, coupled with advanced whole-plant imaging methods, we further separate electric from Ca^2+^/ROS plant-to-plant signals and transcriptional landscapes and show that Ca^2+^/ROS plant-to-plant signals, as well as the function of several EL stress-induced Ca^2+^/ROS-dependent transcripts, are required for EL stress acclimation in a group of plants that touch each other. Our study reveals that plants that live together in a community and physically touch each other establish an aboveground signaling network that enhances their collective resilience to stress.

## Results

### Plants that touch each other aboveground are more resilient to stress

Plants growing within many ecosystems in nature physically touch each other, and this physical touch can mediate electric and/or Ca^2+^/ROS P-T-P signaling between them (*9*, *17*). We therefore hypothesized that plants living in a group or a community, touching each other, will exchange P-T-P signals with each other, and that this signaling process will allow the entire community to become more resilient to stress. To test this hypothesis, we compared the transcriptome and stress resilience of individual *Arabidopsis thaliana* (*A. thaliana*) plants to that of a group of *A. thaliana* plants that physically touch each other (Fig. 1A; SI Appendix, Fig. S1). We found that, compared to individual plants, the transcriptome of plants that physically touch each other for 1 hour or 1 week contained a significantly higher number of stress and hormone response transcripts (2,248 and 1,780 transcripts significantly altered in response to a 1 hour or 1 week of touching, respectively, with 595 transcripts common to these two responses; Figs. 1B, 1C; Datasets S1-S2). These were especially enriched in touch, cold, waterlogging, ROS, and salicylic acid (SA) response transcripts, as well as included transcripts involved in responses to EL stress, salt, wounding, and brassinosteroids (Fig. 1D; Dataset S3). In agreement with these findings, compared to plants living in a group and touching each other, individual plants accumulated more anthocyanins (a common sign of stress in *A. thaliana*), as well as experienced more cellular damage (measured by ion leakage from injured cells), when challenged by growth under EL stress for 24 hours (Fig. 1E; SI Appendix, Fig. S1). The findings presented in Fig. 1 reveal that *A. thaliana* plants that physically touch each other are primed to resist stress and can withstand EL stress better than individual plants that do not touch each other.

**Fig. 1.**
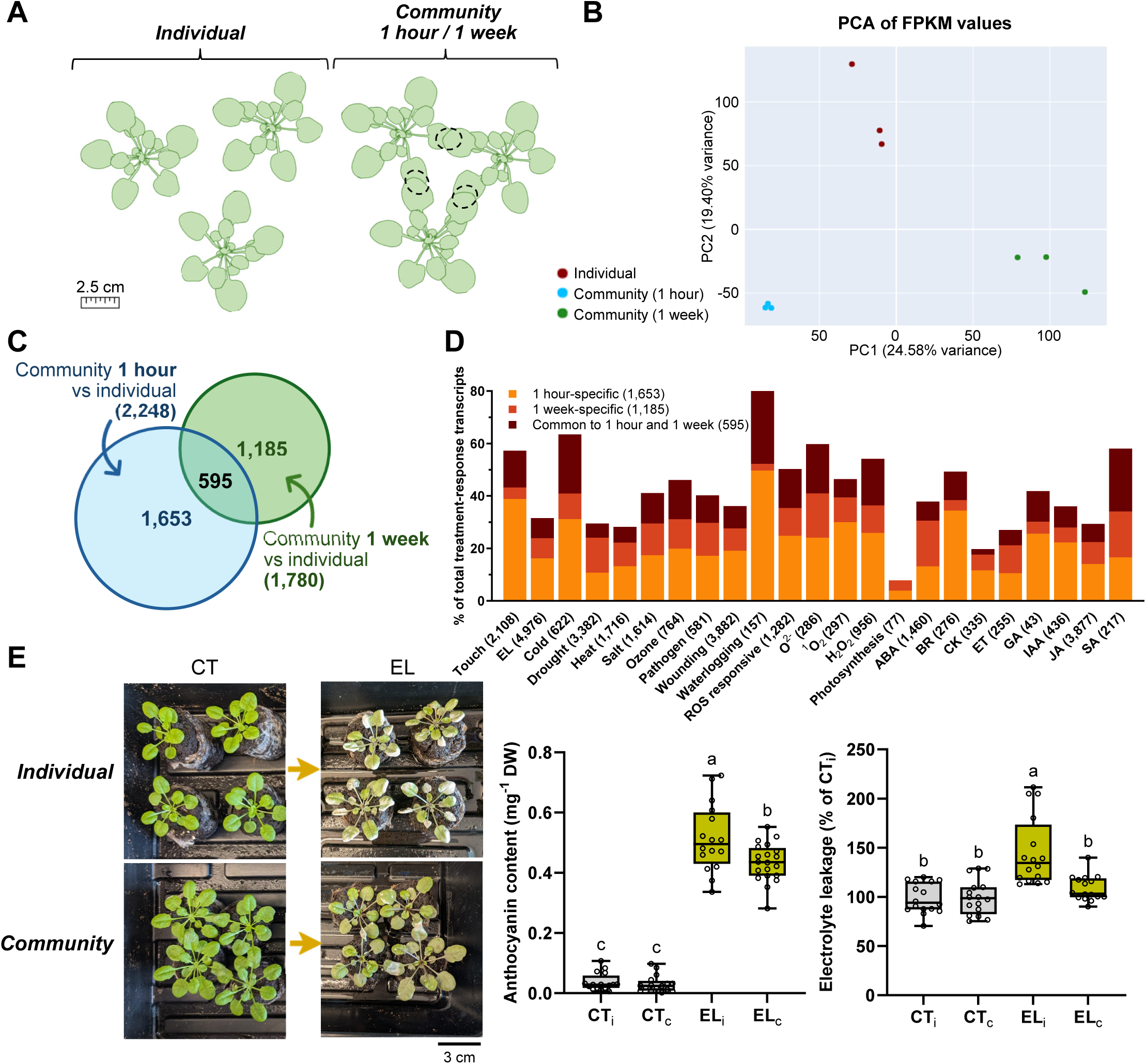
Plants that touch each other aboveground are more resilient to excess light stress. A. The experimental setup used to study plants (*Arabidopsis thaliana*) growing individually or in a community touching each other aboveground (SI Appendix, Fig. S1). B. A PCA plot showing the distinct transcriptomics grouping of plants growing individually or in a community touching each other for 1 hour or 1 week (*N*=3). C. A Venn diagram showing the overlap in transcripts significantly altered in their expression in plants growing as a community and touching each other for 1 hour or 1 week, compared to plants growing individually. D. Representation of different stress-, hormone-, and reactive oxygen species (ROS)-response transcripts among transcripts significantly altered in their expression in plants living as a community and touching each other compared to individual plants (from C). E. Decreased levels of anthocyanins and decreased cellular damage (measured as ion leakage from cells) in plants living as a community touching each other, compared to individual plants, subjected to excess light stress for 24 hours. A representative image of the experimental setup is shown on left, decreased accumulation of anthocyanins (*N*=16) is shown in the middle, and decrease cell death (*N*=16) is shown on right. Significance for B and C was determined by negative binomial Wald test followed by Benjamini–Hochberg correction (*N*=3), and significance for E was determined by one-way ANOVA followed by a Fisher’s LSD post hoc test (different letters denote statistical significance at p ≤ 0.05). Abbreviations: ABA, abscisic acid; BR, brassinolide; CK, cytokinins; CT_i_, individual plants under control conditions; CT_c_, plants growing as a community under control conditions; EL, excess light; EL_i_, individual plants subjected to excess light; EL_c_, plants growing as a community subjected to excess light; ET, ethylene; GA, gibberellic acid; IAA, indole-3-acetic acid; JA, jasmonic acid; SA, salicylic acid.

### Separating EL stress-induced P-T-P electric and Ca^2+^/ROS signals

Two or more plants that physically touch each other aboveground, via leaf-to-leaf contacts, were found to transfer electric and ROS signals between each other (*9*). However, whether either or both signals are required to induce resilience in a group of plants that live together (Fig. 1) remains to be answered. To address this question, we studied the transfer of P-T-P membrane potential (electric), and Ca^2+^/ROS signals between 3 different plants that touch each other in a successive arrangement (*i.e.,* transmitter - T, mediator - M, and receiver - R) in response to EL stress applied using a fiber optic to a single leaf of the transmitter plant (a leaf not involved in the P-T-P connection; termed the local leaf; L; Fig. 2; SI Appendix, Fig. S2). To separate the P-T-P electric signal from the P-T-P Ca^2+^/ROS signal we either used wild type (WT) or a mutant that transmits systemic electric but not Ca^2+^/ROS signals [*HYDROGEN PEROXIDE INDUCED Ca^2+^ INCREASES 1*; *hpca1*(*18*–*20*)], as the mediator plant (Fig. 2; SI Appendix, Fig. S2). This mutant was previously found to transmit the electric signal between an EL stress-treated local leaf and multiple systemic leaves within a plant, without transmitting the Ca^2+^/ROS signal (*18*). As shown in Fig. 2, the application of EL stress to a local leaf of the transmitter plant in the WT-WT-WT arrangement resulted in the transfer of P-T-P electric and Ca^2+^/ROS signals from the transmitter plant, through the mediator plant, to the receiver plant (no P-T-P ROS signals were detected in similarly arranged plants, not subjected to the EL stress; SI Appendix, Fig. S3). In contrast, in the WT-*hpca1*-WT setting, only the electric signal was transferred all the way from the transmitter to the receiver plant, while the P-T-P Ca^2+^/ROS signal was restricted to the transmitter plant (Fig. 2). The two arrangements shown in Fig. 2 (*i.e,* WT-WT-WT and WT-*hpca1*-WT) allowed us therefore to separate the electric from the Ca^2+^/ROS P-T-P signals during EL-driven P-T-P signaling.

**Fig. 2.**
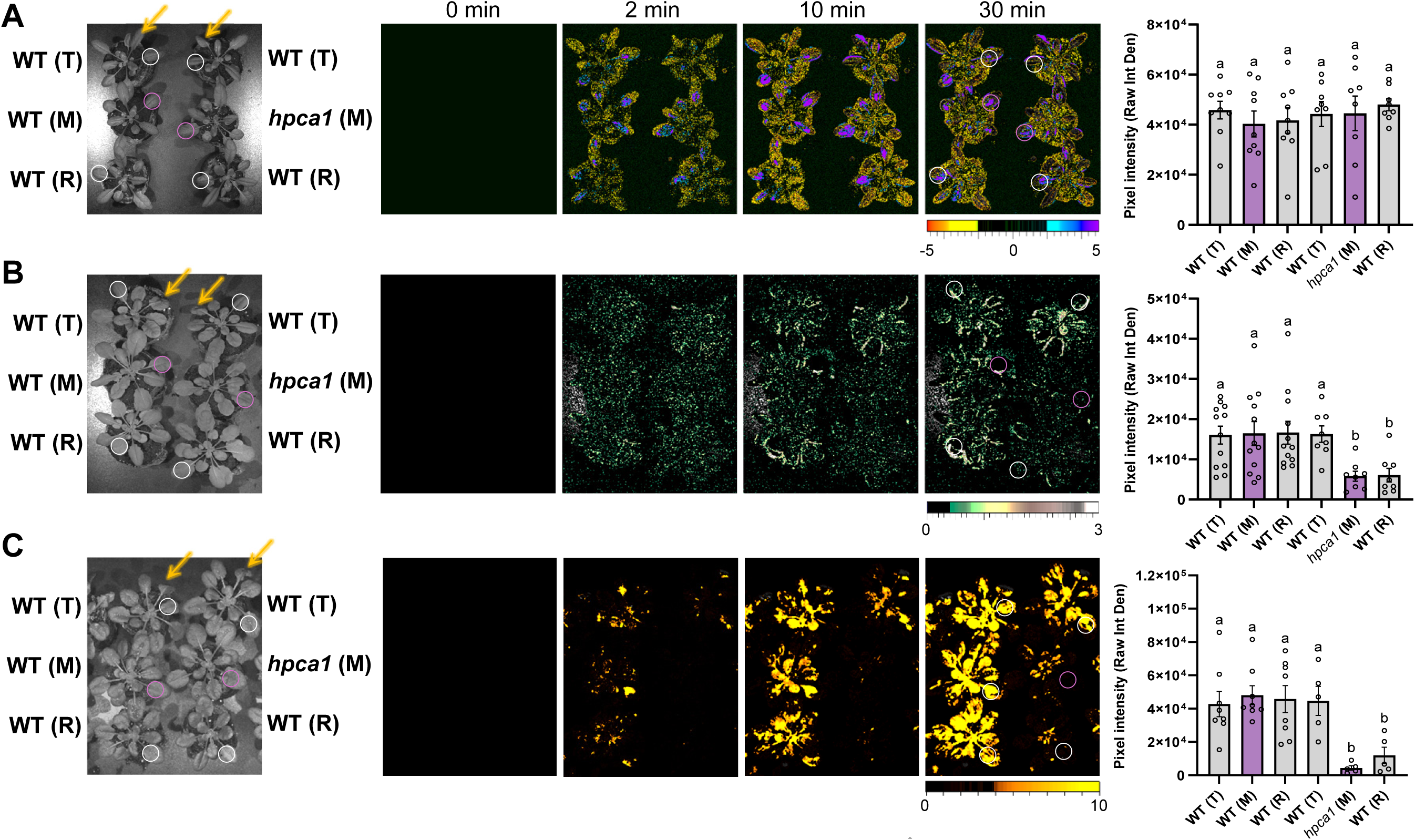
Plants that touch each other aboveground exchange plant-to-plant electric and Ca^2+^/ROS signals in response to stress. A. A bright field image showing the experimental setups used to study the transfer of the plant-to-plant (P-T-P) membrane potential (electric) signal between three successively arranged plants touching each other (transmitter – T, mediator – M, and receiver – R; Left; SI Appendix, Fig. S2), time lapse images of membrane potential change in plants touching each other in response to an excess light (EL) stress applied to one leaf of the transmitter plant (Middle), and a bar graph showing statistical analysis of membrane potential change intensity at 30 min (Right; *N*=8). B. Same as A, but for the P-T-P calcium (Ca^2+^) signal (*N*=8). C. Same as A, but for the P-T-P reactive oxygen species (ROS) signal (*N*=5). See SI Appendix, Fig. S3 for imaging of plants touching each other but not subjected to EL stress. Local leaf subjected to EL stress is indicated with an arrow. White circles represent regions of interest used for signal intensity measurements. Significance was determined by a one-way ANOVA followed by a Fisher’s LSD post hoc test (different letters denote statistical significance at p ≤ 0.05). Abbreviations: HPCA1, HYDROGEN PEROXIDE INDUCED Ca^2+^ INCREASES 1; M, mediator; R, receiver; T, transmitter; WT, wild type.

### Plant-to-plant Ca^2+^/ROS signals are required for enhanced acclimation of receiver plants to EL stress

Using the two setups shown in Fig. 2, we asked what P-T-P signals are required to induce acclimation to EL in the receiver plant in response to an EL stress applied to the transmitting plant? In agreement with previous studies of individual plant acclimation (*21*–*23*), when EL stress was applied to the local (L) leaf of the transmitting plant, it induced acclimation to EL stress in the systemic leaf of the same plant (leaf S1; Fig. 3; SI Appendix, Fig. S4A). Remarkably, in the WT-WT-WT setup, it also induced acclimation in a systemic leaf of the receiver plant (leaf S3; Fig. 3; in agreement with Fig. 1). In contrast, while application of EL to the local leaf of the transmitter plant in the WT-*hpca1*-WT setup induced acclimation in the systemic leaf of this plant (S1), it was unable to induce acclimation in the systemic leaf of the receiver plant (leaf S3; Fig. 3). This finding revealed that, in the absence of the P-T-P Ca^2+^/ROS signal, the transfer of the P-T-P electric signal between all 3 plants in the WT-*hpca1*-WT setup (Fig. 2) is insufficient to induce acclimation to EL in the receiver plant (Fig. 3). The transfer of the electric and Ca^2+^/ROS, or just the Ca^2+^/ROS, P-T-P signal(s) is therefore likely required for the acclimation of a community or a group of plants to EL stress (Figs. 1-3; SI Appendix, Figs. S2 and S4A).

**Fig. 3.**
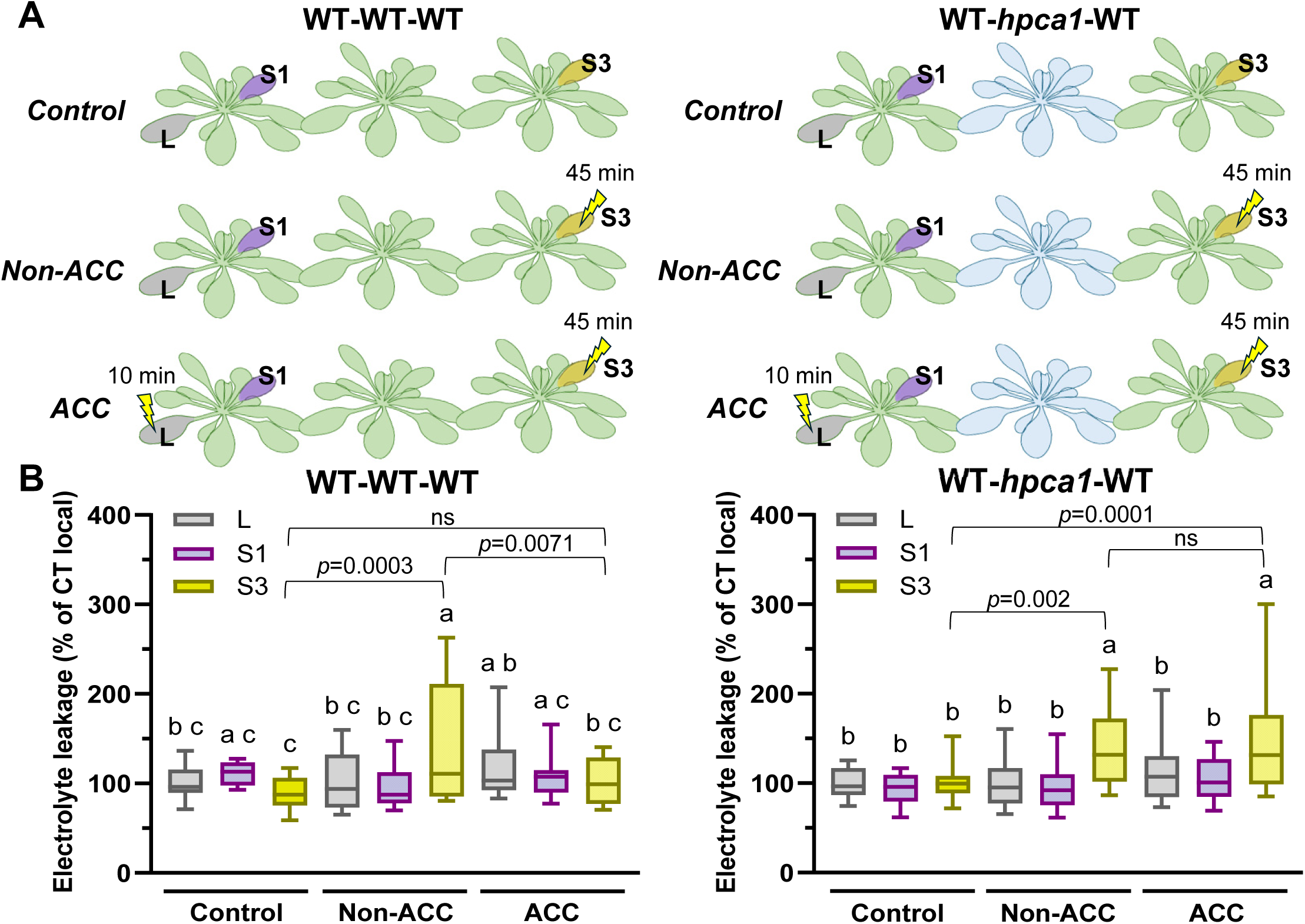
Plant-to-plant Ca^2+^/ROS signals are required for plant acclimation to excess light stress. A. The experimental setup used to study local (L) and systemic (S1, S3) leaf acclimation to excess light (EL) stress in three successively arranged plants touching each other (SI Appendix, Fig. S4A). B. Bar graphs showing cellular damage, measured as ion leakage from cells, in leaves L, S1 and S3 following an extended EL stress (45 min) in the presence or absence of a short (10 min) EL stress applied to leaf L of the transmitter plant (Bottom; N=7). Significance was determined by a one-way ANOVA followed by a Fisher’s LSD post hoc test comparing L, S1 and S3 of non-acclimated (ACC) and ACC plants versus control plants (different letters denote statistical significance at p ≤ 0.05). Abbreviations: ACC, acclimated; HPCA1, HYDROGEN PEROXIDE INDUCED Ca^2+^ INCREASES 1; ns, non-significant; WT, wild type.

### Transcriptomics analysis of P-T-P signaling in response to EL stress

To determine what transcriptomic responses are associated with the electric or Ca^2+^/ROS P-T-P signals (Fig. 2), we conducted an RNA-Seq analysis of the local (L) and systemic (S1-S3) leaves of plants arranged in the WT-WT-WT and WT-*hpca1*-WT setups in response to an EL stress applied to a single leaf of the transmitter plant (not involved in the P-T-P interaction; Fig. 4A, SI Appendix Figs. S2 and S5; Dataset S4-S10). Plants, arranged in rows of 3 as described above (Fig. 2), were either treated with EL applied to leaf L of the transmitter plant, or untreated (Control; CT), incubated for 30 min and sampled (Fig. 4A). Local (L) and systemic (S1-S3) leaves of EL stress-treated and untreated setups were pooled from 60 plants (belonging to 20 different rows) and used in 3 biological repeats for RNA-Seq analysis. Transcripts significantly altered in their expression in L, S1, S2, S3 leaves of treated and untreated plants were determined (Fig. 4B). Once significant transcripts were identified in each setup type, the overlap between transcripts significantly altered in the WT-WT-WT and WT-*hpca1*-WT setups was determined (Fig. 4C).

**Fig. 4.**
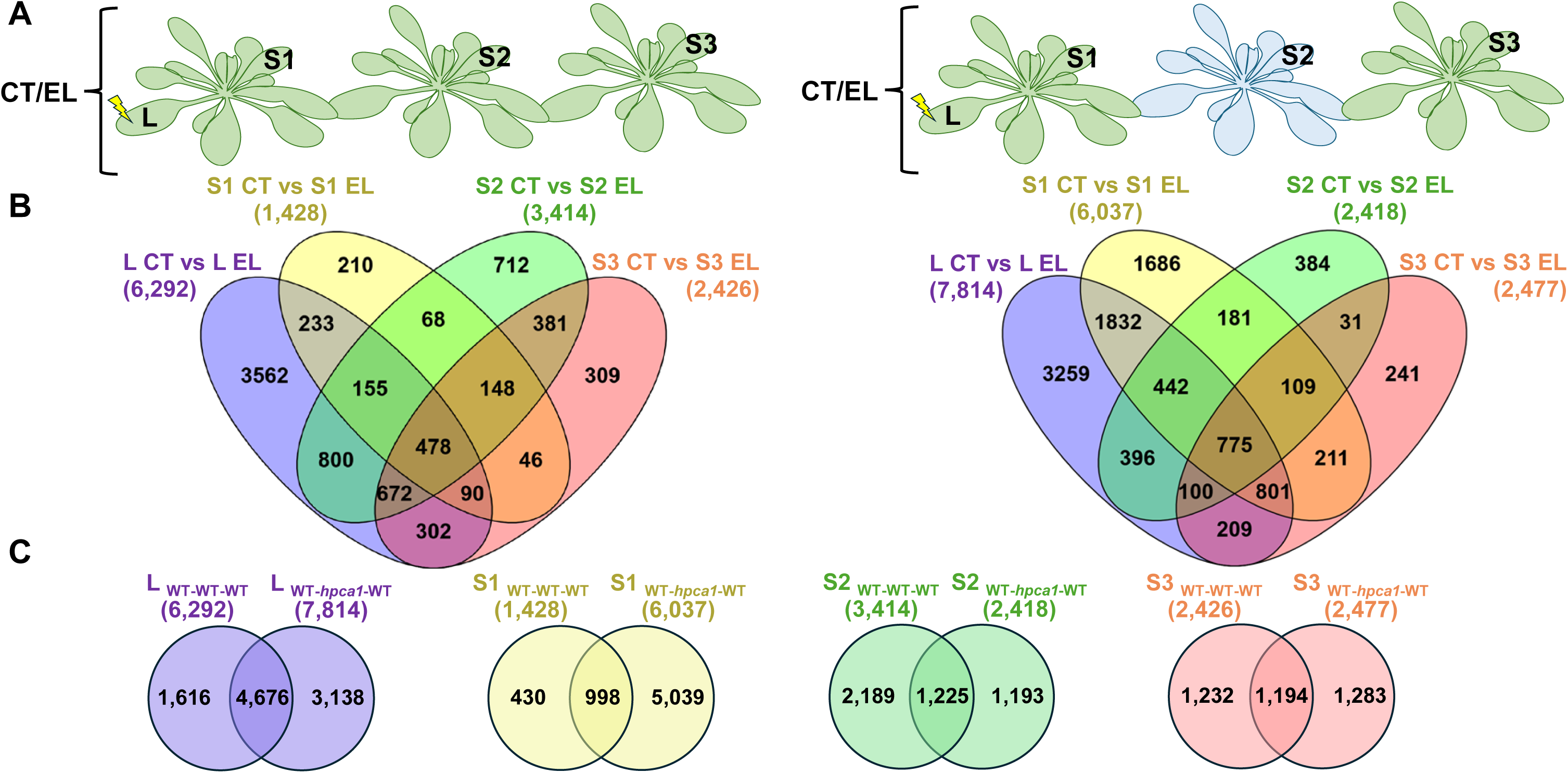
Transcriptomics analysis of plant-to-plant signaling. A. The experimental setups used for the transcriptomics study of plant-to-plant (P-T-P) signaling between three successively arranged plants touching each other (transmitter – T, mediator – M, and receiver – R). B. Venn diagrams showing the overlap between transcripts significantly altered in their expression in local (L) and systemic (S1-S3) leaves of plants from the two setups shown in A, in response to excess light (EL) stress applied to leaf L (see SI Appendix, Fig. S2). C. Venn diagrams showing the overlap between L and S1-S3 from the two different setups shown in A and B. Significance was determined by negative binomial Wald test followed by Benjamini–Hochberg correction (*N*=3). Abbreviations: CT, control; EL, excess light; HPCA1, HYDROGEN PEROXIDE INDUCED Ca^2+^ INCREASES 1; L, local; S1, systemic leaf 1; S2, systemic leaf 2; S3, systemic leaf 3; WT, wild type.

While the steady-state level of over 6,000 transcripts was significantly altered in the local leaves of plants from both setups (WT-WT-WT and WT-*hpca1*-WT), in response to EL stress applied to the L leaf of each treated setup (Fig. 4B; with an overlap of over 4,500 transcripts; Fig. 4C), except for S1 in the WT-*hpca1*-WT setup, fewer transcripts (ranging from 1,400 to 3,400) were altered in their expression in the systemic leaves of each setup in response to EL stress applied to the local leaf (leaves S1-S3; Fig. 4B; Datasets S5-S6; SI Appendix, Figs. S6-S7). Notably, transcripts present in S2 and S3 in both setups had a considerable overlap with their corresponding local leaf (1,500-2,100 in the WT-WT-WT setup and 1,700-1,800 in the WT-*hpca1*-WT setup), as well as an overlap of about 1,200 transcripts with each other (Figs. 4B, 4C; Datasets S5-S6). These findings suggest that P-T-P signaling (Fig. 2) can result in the altered expression of thousands of transcripts in the systemic leaves of mediator and receiver plants (leaves S2 and S3) that are connected to the transmitter (leaves L, S1) plant through P-T-P leaf connections, but are not directly exposed to the EL stress (only applied using a fiber optic to leaf L of the transmitter plant; Figs. 2, 3, 4A; SI Appendix, Fig. S2). As many of the transcripts altered in the systemic leaves of the mediator and receiver plants (leaves S2 and S3 in both setups) had a considerable overlap with the local (L) leaf of the transmitter plant that was exposed to the EL stress (Figs. 4B, 4C), it is likely that the transfer of some of the different P-T-P signals between the successively arranged plants that touched each other had an important biological context and could induce acclimation to EL stress (Fig. 3).

The exceptionally large number of transcripts altered in the systemic leaf of the transmitter plant (6,037; leaf S1; Fig. 4B) in the WT-*hpca1*-WT setup, compared to the low number of overall transcripts in the S1 leaf in the WT-WT-WT setup (1,428; Fig. 4B), could suggest that the disruption in P-T-P signaling caused by the inclusion of the *hpca1* mutant as the mediator plant in the WT-*hpca1*-WT setup (Fig. 2) caused some type of a feedback response in the transmitter plant of this setup. These transcripts were enriched in SA, H_2_O_2_, ozone and ROS response transcripts (Dataset S7), suggesting that the inability of the transmitter plant to induce the Ca^2+^/ROS signals in the mediator (*hpca1*) plant in the WT-*hpca1*-WT setup could have resulted in a potential Ca^2+^/ROS-associated feedback response.

### Identification of putative electric- and Ca^2+^/ROS-associated P-T-P signaling transcripts

Compared to the WT-WT-WT setup, the Ca^2+^/ROS P-T-P signal was suppressed in mediator and receiver plants belonging to the WT-*hpca1*-WT setup (Fig. 2), in which the systemic leaf of the receiver plant (leaf S3) was unable to acclimate to EL stress (Fig. 3). In contrast, the electric P-T-P signal was detected in all plants belonging to both setups (Fig. 2). This difference allowed us to separate transcriptomics responses to the electric P-T-P signal from those to the Ca^2+^/ROS P-T-P signal (Fig. 5). Thus, we identified transcripts absent from leaves S2 and S3 in the WT-*hpca1*-WT setup (compared to the WT-WT-WT setup; marked with an ‘X’ in Fig. 5; 37 transcripts; Dataset S8) and termed them ‘putative Ca^2+^/ROS-dependent transcripts’, as well as identified transcripts present in leaves S2 and S3 in the WT-*hpca1*-WT setup (compared to the WT-WT-WT setup; marked with an ‘√’ in Fig. 5; 589 transcripts; Dataset S9) and termed them ‘putative electric signal-dependent transcripts’.

**Fig. 5.**
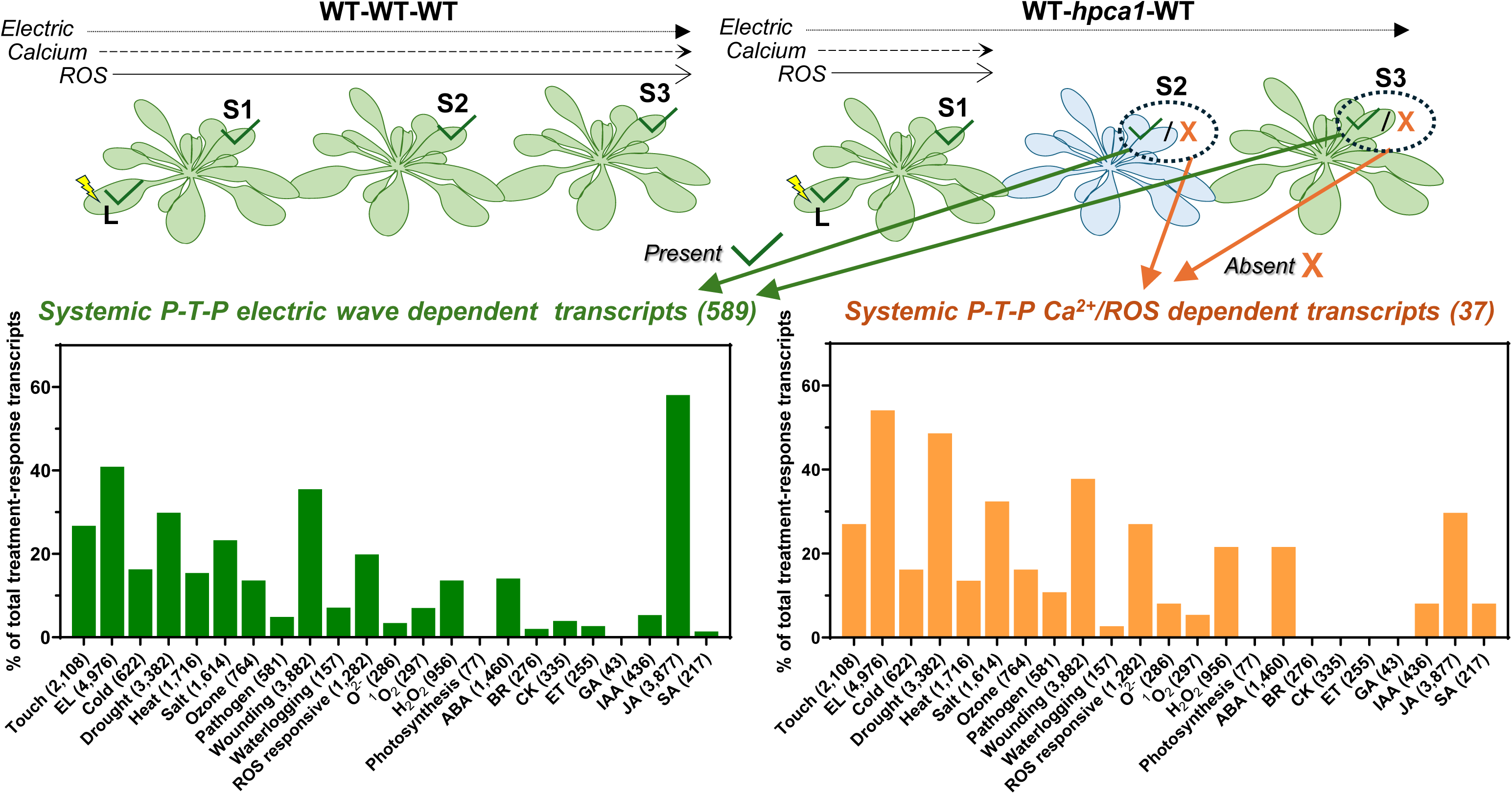
Identification of putative electric and Ca^2+^/ROS associated transcripts significantly altered in their expression during plant-to-plant signaling. The scheme used to identify putative plant-to-plant (P-T-P) electric and Ca^2+^/ROS associated transcripts significantly altered in their expression during excess light (EL) stress driven P-T-P signaling (Top), and representation of different stress, hormone, or ROS response transcripts among the two different groups (putative electric-dependent – 589 transcripts, and putative Ca^2+^/ROS-dependent – 37 transcripts; Bottom). Significance was determined by negative binomial Wald test followed by Benjamini–Hochberg correction (*N*=3). Abbreviations: ABA, abscisic acid; BR, brassinolide; CK, cytokinins; EL, excess light; ET, ethylene; GA, gibberellic acid; HPCA1, HYDROGEN PEROXIDE INDUCED Ca^2+^ INCREASES 1; IAA, indole-3-acetic acid; JA, jasmonic acid; L, local; P-T-P, plant-to-plant; ROS, reactive oxygen species; S1, systemic leaf 1; S2, systemic leaf 2; S3, systemic leaf 3; SA, salicylic acid; WT, wild type.

While wounding and touch response transcripts were equally represented in both groups of transcripts, putative electric signal-dependent transcripts were especially enriched in jasmonic acid (JA) response transcripts, while putative Ca^2+^/ROS-dependent transcripts were especially enriched in EL stress, drought, salt, abscisic acid (ABA), and H_2_O_2_ response transcripts (Fig. 5; Dataset S10). As P-T-P Ca^2+^/ROS signaling was required for receiver plant acclimation to EL stress (Fig. 3), we hypothesized that some of the putative P-T-P Ca^2+^/ROS-response transcripts present in the WT-WT-WT setup in leaves S2 and S3, but absent in the WT-*hpca1*-WT setup (37 transcripts; Fig. 5 and SI Appendix, Fig. S8; Dataset S8), are required for this acclimation process.

### Genes encoding putative P-T-P Ca^2+^/ROS-response signaling transcripts are essential for plant acclimation to EL stress

To test the hypothesis that putative P-T-P Ca^2+^/ROS-response transcripts are required for plant acclimation to EL stress, we obtained mutants deficient in the expression of several of these transcripts and tested their ability to acclimate to EL stress (Fig. 6). We define acclimation to EL stress as suppressed levels of leaf injury (measured by low levels of ion leakage) in a local or systemic leaf that was pre-treated with a short period of EL stress (usually 10 min), allowed to acclimate (usually for 45-60 min), and challenged by a long period of EL stress of 45-60 min (compared to a local or systemic leaf that was not pre-treated with the short EL stress prior to being exposed to a long period of EL stress and displays a high level of ion leakage following the extended EL stress period; SI Appendix, Fig. S4B; (*21*–*23*). Altogether, we studied the role of 6 different genes (using two independent mutant alleles per locus; Dataset S11), identified as encoding Ca^2+^/ROS-response transcripts to EL stress (Fig. 6; SI Appendix, Fig. S8; Dataset S8) and identified 5 (*i.e.,* AT2G38470, AT1G33760, AT1G30730, AT1G51800, AT5G65300) that had altered ability to acclimate to EL stress (Fig. 6). These transcripts were chosen for analysis based on their expression pattern and putative function (Figs. 4, 5; SI Appendix, Fig. S8; Dataset 11).

**Fig. 6.**
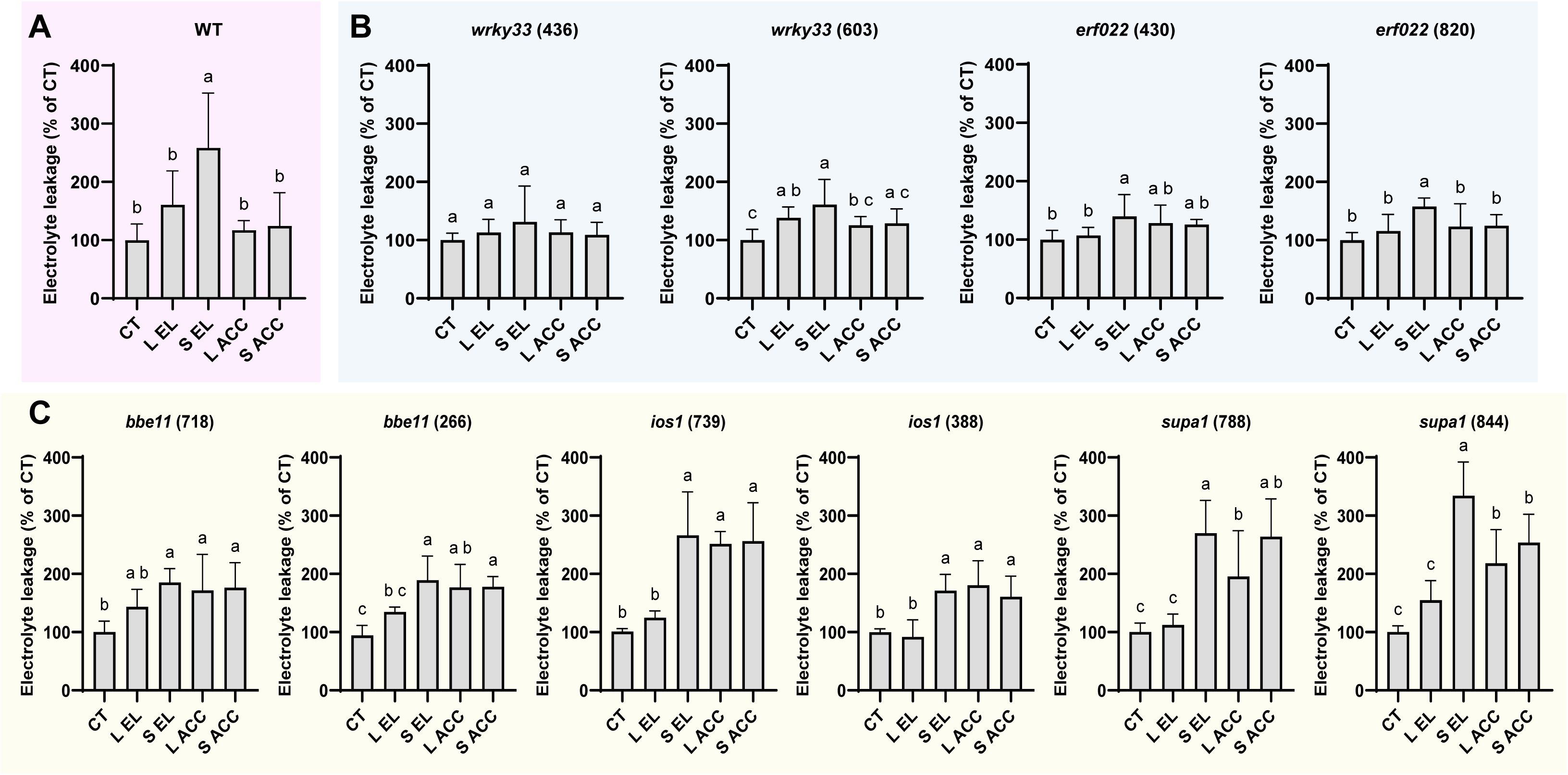
Genes encoding Ca^2+^/ROS associated plant-to-plant signaling transcripts are essential for plant acclimation to excess light stress. A. Decrease in cellular damage (measured as ion leakage) in local (L) or systemic (S) leaves from wild type plants subjected to a prolonged (45 min) excess light (EL) stress following a short (10 min) pretreatment of EL stress applied to leaf L (*i.e.,* acclimation to EL stress; SI Appendix, Fig. S4B; *N*=5). B. Enhanced basal acclimation to EL stress in mutants deficient in the expression of different Ca^2+^/ROS associated plant-to-plant (P-T-P) signaling transcripts (*N*=4). C. Failure of mutants deficient in the expression of different Ca^2+^/ROS associated P-T-P signaling transcripts to acclimate to EL stress following a short (10 min) pretreatment of EL stress applied to leaf L (*N*=4). Significance was determined by a one-way ANOVA followed by Fisher’s LSD post hoc test (different letters denote statistical significance at p ≤ 0.05). Abbreviations: AAC, acclimation; BBE, berberine bridge enzyme; CT, control; EL, excess light; ERF, ethylene response factor; IOS, impaired oomycete susceptibility; L, local; S, systemic; SUPA, salt-up a; WRKY, WRKY DNA- binding transcriptional regulator; WT, wild type.

While WT plants were able to locally and systemically acclimate to an extended (45 min) EL stress following a short (10 min) pre-treatment of EL stress applied to a single leaf [leaf L; Fig. 6A and SI Appendix, Fig. S4B; previously reported by (*21*–*23*)], two independent mutant alleles per locus of *WRKY DNA-BINDING PROTEIN 33* (*WRKY33*; AT2G38470; *24*), a transcriptional regulator involved in responses to various biotic and abiotic stresses, and *ETHYLENE RESPONSE FACTOR 22* (*ERF22*; AT1G33760; *25*), a transcriptional regulator involved in ethylene responses, appeared to be constitutively resilient to EL stress and did not display enhanced ion leakage in the presence or absence of the pre-treatment (Fig. 6B). These findings suggest that the products of these transcripts could function as suppressors of the EL stress acclimation response and that in their absence the EL acclimation response was constitutively turned ‘on’. In contrast, two independent mutant alleles per locus for *BERBERINE BRIDGE ENZYME 11* (*BBE11*; AT1G30730; *26*), an oligogalacturonide oxidase involved in damage-associated molecular patterns (DAMPs) signaling as suppressor of immune responses, *IMPAIRED OOMYCETE SUSCEPTIBILITY 1* (*IOS1*; AT1G51800; *27*), an LRR-RLK protein family member involved in Pattern-Triggered Immunity activation, and *SALT-UPA 1* (*SUPA1*; AT5G65300; *28*), a peroxisomal protein of unknown function, rapidly expressed in response to different biotic and abiotic stresses, were unable to systemically or locally acclimate to an extended EL stress following a short local application of EL stress. Thus, two independent mutants deficient in the expression of each of these transcripts displayed enhanced local and systemic leaf ion leakage in response to EL stress in the presence or absence of the EL stress pre-treatment applied to the local leaf (Fig. 6C). These findings suggested that, when these transcripts are dysregulated, plants cannot activate the mechanisms required for EL stress acclimation and cannot prevent the damage caused to leaves by this stress (Fig. 6C).

The findings presented in Figs. 2-6 suggest therefore that several different transcripts associated with the P-T-P Ca^2+^/ROS signal are required for plant acclimation to EL stress. In addition, they demonstrate that acclimation to EL stress and responses to biotic challenges may share similar regulators and/or pathways.

### Impaired accumulation of ROS in *bbe11* and *ios1* mutants

The failure to acclimate to EL stress (Fig. 6), and the link to Ca^2+^/ROS signaling (Fig. 5; SI Appendix, Fig. S8), of some of the Ca^2+^/ROS-associated P-T-P signaling mutants/transcripts, could suggest that they are deficient in intraplant systemic ROS signaling (*i.e.,* from their local to systemic leaves). Such deficiency could explain why they are unable to acclimate (Fig. 6C), as well as their association with P-T-P Ca^2+^/ROS signaling (Fig. 5; SI Appendix, Fig. S8), that requires each plant in the path of the P-T-P signal to mediate intraplant systemic ROS signaling (Fig. 2). We therefore tested whether mutants deficient in the expression of two of these transcripts (*i.e., ios1* and *bbe11*; Fig. 6C) can mount a systemic whole plant ROS response following application of EL to their local leaf. As shown in Fig. 7, two independent mutant alleles for *ios1* or *bbe11* were unable to mount a systemic ROS response to a local EL stress. This finding provides a potential mechanistic explanation for the role of some of the Ca^2+^/ROS transcripts during P-T-P signaling. They could be required for transmitting the intraplant systemic ROS signal from the local to the systemic leaves of each plant, and hence required for the transfer a P-T-P signal through each plant from its local leaf that touches another plant, or is stressed, to its systemic leaves that touch the next, or other plants in the community.

**Fig. 7.**
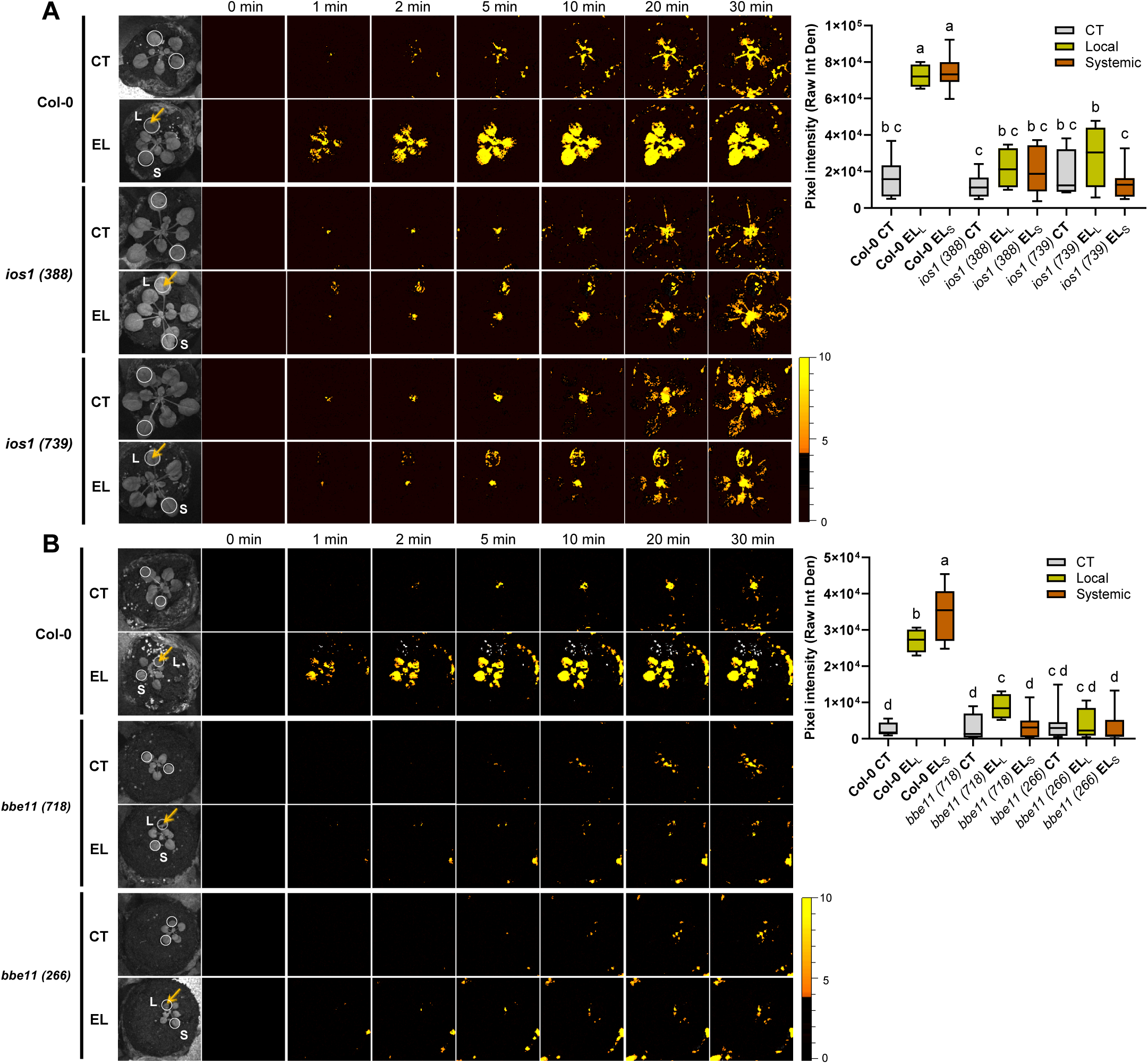
Impaired accumulation of ROS in two Ca^2+^/ROS associated plant-to-plant signaling mutants that are unable to acclimate to excess light stress. A. Time lapse images of whole plant ROS accumulation in response to an excess light (EL) stress applied to one leaf (indicated with an arrow on the bright field image) of a wild type (Col-0) or two independent alleles for the *ios1*mutation (Left), and a bar graph showing statistical analysis of ROS accumulation intensity at 30 min (Right; *N*=4). White circles represent regions of interest used for signal intensity measurements. B. Same as A, but for wild type and two independent alleles for the *bbe1* mutation (*N*=4). Significance was determined by a one-way ANOVA followed by a Fisher’s LSD post hoc test (different letters denote statistical significance at p ≤ 0.05). Abbreviations: bbe, berberine bridge enzyme; CT, control; Col-0, Columbia Col-0; EL, excess light; ios, impaired oomycete susceptibility; L, local; S, systemic.

### Gene regulatory network analysis of the P-T-P signaling response

To determine what pathways and networks control, are controlled by, or are associated with, P-T-P signaling, we conducted a gene regulatory network (GRN) analysis of the transcriptomic response of leaf S3 of receiver plants, following the application of EL stress to leaf L of transmitter plants, in the WT-WT-WT and WT-*hpca1*-WT setups (Datasets S12-S13). As shown in Fig. 8A and Dataset S12, the pathways and networks unique to leaf S3 of receiver plants in the WT-WT-WT setup included flavonoid biosynthesis and glutathione metabolism. As both flavonoids and glutathione are known to play a key role in ROS metabolism (*29*–*31*), the absence of these networks from leaf S3 in the WT-*hpca1*-WT setup is in agreement with our findings that the P-T-P ROS signal did not reach this leaf (Fig. 2), and that this leaf was unable to acclimate to EL stress (Fig. 3), in the WT-*hpca1*-WT setup. Among the pathways and networks unique to leaf S3 of the WT-*hpca1*-WT setup were several stress and insect/wounding associated pathways (Fig. 8A, Dataset S13). As responses to wounding and insect attack are associated with electric signals and the plant hormone JA (*32*, *33*), these networks could be associated with the electric P-T-P signal that reached leaf S3 of receiver plants, in the absence of the ROS P-T-P signal, in the WT-*hpca1*-WT setup (Fig. 2). In addition to the broad GRN analyses shown in Fig. 8A, we also conducted a more specific GRN analysis focused on IOS1- and BBE11- associated networks of leaf S3 in the WT-WT-WT setup (Datasets S14-S15). As shown in Fig. 8B, IOS1 appeared to be under the control of multiple TFs involved in flowering, stress/drought response, circadian rhythm, and chlorophyll biosynthesis. Interestingly, included within these TFs was ERF22 that was also found to have an impaired EL stress acclimation (Fig. 6B). As shown in Fig. 8C, BBE11 appeared to be under the control of multiple TFs involved in leaf and flower development, salt/ABA responses, photosynthesis, and circadian rhythm. Taken together, our transcriptomics and GRN analyses reveal that P-T-P signaling could transmit, or be associated with, a broad spectrum of signals involved in multiple processes and pathways (Figs. 1, 4, 5, 8; Datasets S12-S15).

**Fig. 8.**
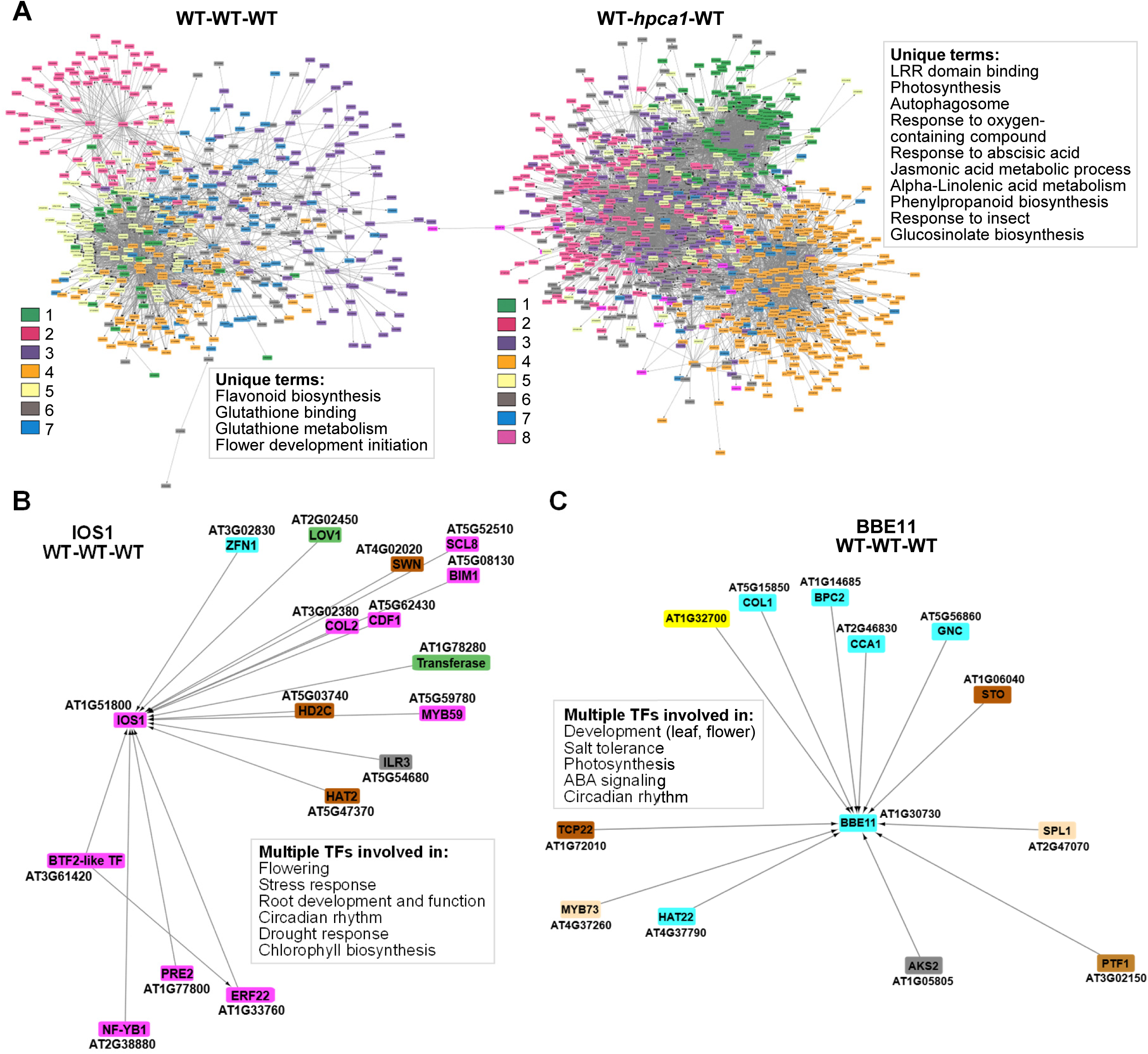
Gene regulatory network analysis of the plant-to-plant signaling response. A. Gene regulatory network (GRN) analysis of the transcriptomic response of leaf S3 of receiver plants, following the application of excess light (EL) stress to the local leaf of transmitter plants, in the WT-WT-WT (left) and WT-*hpca1*-WT setups (right). B. Same as in A, but for the specific clusters involving IOS1 (left) and BBE11 (right) in leaf S3 of the WT-WT-WT setup. The composition, abbreviations, and gene ontology annotation of all clusters shown in A and B are described in Datasets S12-S15.

### Plants that touch each other aboveground have higher basal levels of ROS that can be further elevated upon stress application

As plants touching each other express many stress-response transcripts, even in the absence of stress (compared to plants growing individually; Fig. 1), we hypothesized that they would also have a high basal level of ROS that could cause them to be more resilient to stress. Testing this hypothesis, we found that *A. thaliana* seedlings growing individually had a lower basal level of ROS, compared to *A. thaliana* seedlings growing in a group, touching each other above- and below-ground (Fig. 9A). Direct H_2_O_2_ measurements (using the Amplex Red method; *18*) in tissues of plants, living alone or in a group, corresponded with our ROS imaging results (Fig. 9B). In agreement with our findings for P-T-P ROS signals in the WT-WT-WT setup (Fig. 2), applying EL stress to several members of a plant group, comprised of multiple *A. thaliana* seedlings growing in soil and touching each other above- and below-ground, resulted in further enhancement in ROS levels in the entire group (Fig. 9C). Thus, much like interactions with a beneficial microbe that causes elevated basal levels of ROS in plants, that could be elevated even further following stress (*34*), plants living together and touching each other displayed a higher basal level of ROS that could be enhanced even further during P-T-P responses to stress (Fig. 9C).

**Fig. 9.**
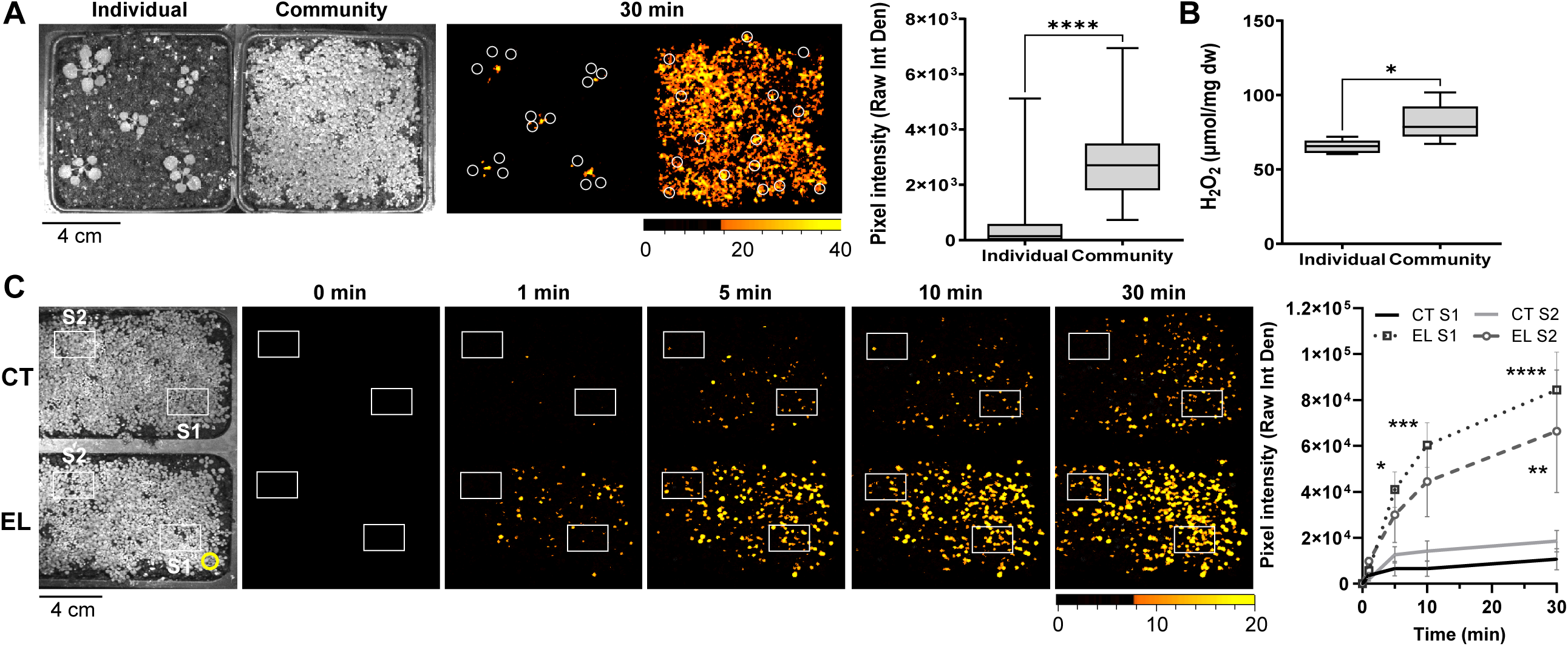
Enhanced basal and stress induced ROS levels in plants that physically touch each other. A. Bright field image (left), reactive oxygen species (ROS) accumulation image (middle), and bar graph showing quantification of the ROS signal at 30 min (*N*=10; right) in single plants (individual) and a dense group of seedlings touching each other below and above ground (community) under control conditions. Significance was determined by Student’s *t* test ***p ≤ 0.001. White circles represent regions of interest used for signal intensity measurements. B. H_2_O_2_ accumulation in leaf tissues of single plants (individual) and a dense group of seedlings touching each other above and below ground (community) under control conditions (Measured using the Amplex Red assay; *18*; *N*=5). Significance was determined by Student’s *t* test with significance levels: *p ≤ 0.05. C. Time lapse images of ROS accumulation in a dense group of seedlings touching each other in response to an EL stress applied to a few plants (yellow circle) and a line graph showing quantification of ROS accumulation in two groups of systemic seedlings (white boxes represent regions of interest used to measure signal intensity) growing 1 (S1) or 8 (S2) cm away from the treated seedlings, respectively (Right; *N*=4). Significance was determined using a two-way ANOVA followed by a Tukey post hoc test (p ≤ 0.05). Abbreviations: CT, control; EL, excess light; ROS, reactive oxygen species.

### H_2_O_2_ accumulation at the leaf-to-leaf contact point between two different plants is required for the transfer of the P-T-P ROS signal between them

Our findings that, in the WT-*hpca1*-WT setup, the Ca^2+^/ROS signal did not transfer through the mediator plant to the receiver plant, while the electric signal did (Fig. 2), suggested that in addition to the electric P-T-P signal, another type of P-T-P signal, related to Ca^2+^/ROS signaling, should be transferred between plants. As H_2_O_2_ can easily diffuse from a plant cell that produces high levels of H_2_O_2_ to a nearby cell that has low levels of H_2_O_2_ (*35*, *36*), and plants that accumulate high levels of ROS in their apoplast could diffuse/secrete H_2_O_2_ into the outer leaf space between two touching leaves, as long as humidity is high, it is possible, in principle, that when a high ROS producing plant physically touches a low ROS producing plant, a similar transfer of H_2_O_2_ will occur from one plant to the other. To test this possibility, we added the H_2_O_2_ scavenging enzyme catalase to the leaf-to-leaf contact points between two different plants and tested whether this addition would suppress the transfer of the P-T-P ROS signal between them. As shown in Figs. 10A, 10B, two wild type plants (a transmitter, T, and a receiver, R), connected via one leaf-to-leaf contact point that included a buffer (indicated with a blue square), were able to transmit the P-T-P ROS signal between them in response to an EL stress applied to one leaf of the transmitter plant (a leaf that was not involved in the P-T-P connection; marked with a yellow arrow). In contrast, when catalase was included in the buffer solution that connected the two plants (indicated with a blue square), the P-T-P ROS signal did not transfer from the transmitter to the receiver plant in response to the EL stress treatment applied to one leaf of the transmitter plant (a leaf that was not involved in the P-T-P connection; marked with a yellow arrow). In support of our findings with catalase (Figs. 10A, 10B), direct measurements of H_2_O_2_ (using the Amplex Red method; *18*), in the buffer placed at the leaf-to-leaf contact point between the transmitter and receiver plants, revealed that H_2_O_2_ accumulated in this buffer in response to EL stress applied to a remote leaf (a leaf that was not involved in the P-T-P connection) of the transmitter plant (Fig. 10C). Taken together, the findings presented in Fig. 10 support a model in which the transfer of the P-T-P ROS signal between two plants that physically touch each other requires the transmitting plant, that accumulates high levels of ROS, to transmit H_2_O_2_ to the receiver plant that did not accumulate high levels of ROS yet. Such transmission could occur via a simple mechanism of diffusion from the apoplast of the high ROS producing plant to the apoplast of the low ROS producing plant, across the leaf-to-leaf touching point, as long as conditions are humid.

**Fig. 10.**
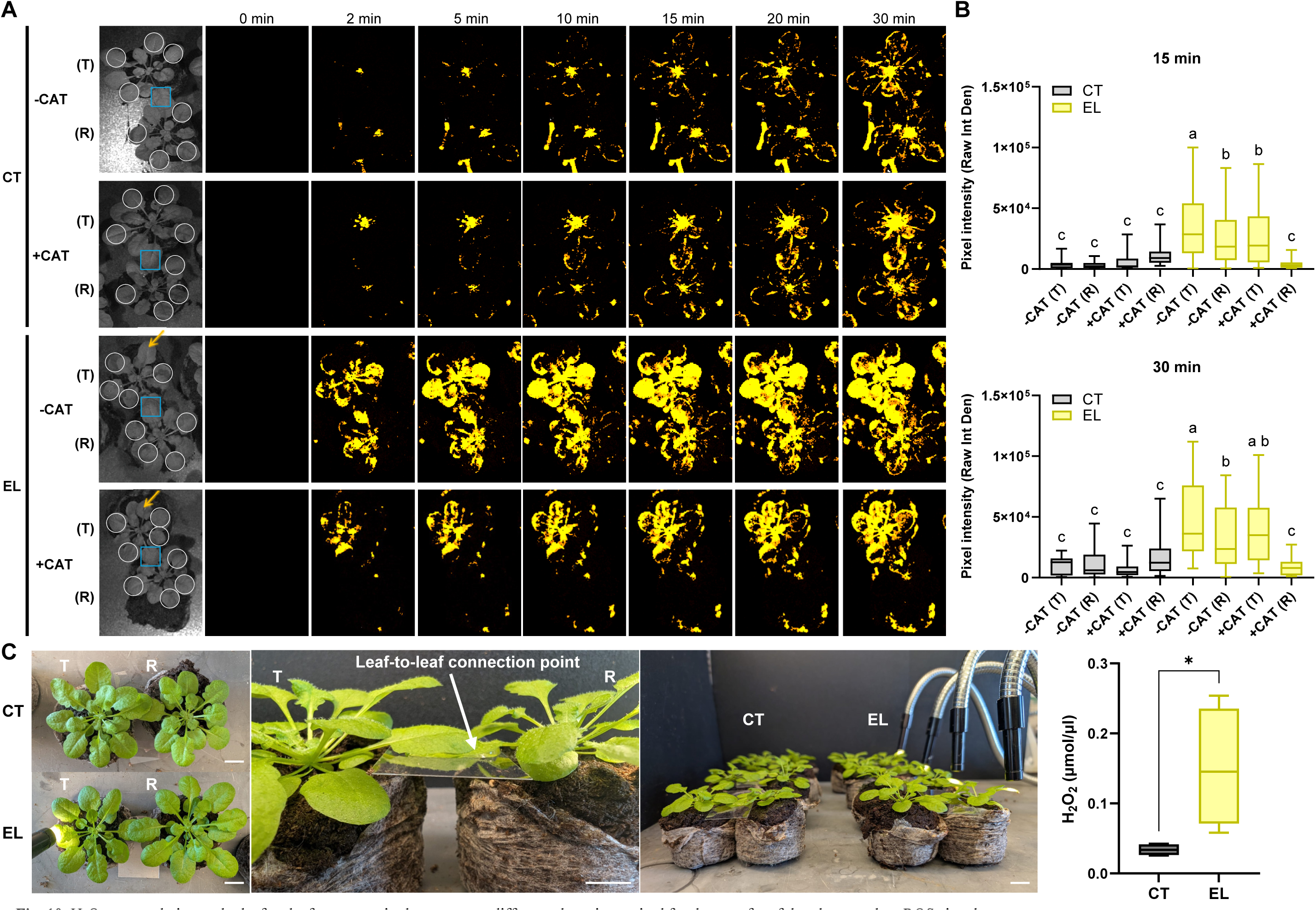
H_2_O_2_ accumulation at the leaf-to-leaf contact point between two different plants is required for the transfer of the plant-to-plant ROS signal. A. Bright field (left) and time lapse images of reactive oxygen species (ROS) accumulation (right), of two plants (T, transmitter, and R, receiver) connected with each other through one leaf-to-leaf contact point (indicated with a blue square) that includes buffer with catalase (+CAT), or just buffer (-CAT). Imaging was conducted following the application of an excess light (EL) stress to a single leaf of the transmitter plant (indicated with an arrow). Regions of interest used to quantify signal intensity are indicated with white circles. B. Bar graphs showing quantification of the ROS signal from A at 15 (top) or 30 (bottom) min following the application of the EL stress to one leaf of the transmitter plant. Significance was determined by one-way ANOVA followed by Fisher’s LSD post hoc test (different letters denote statistical significance at p ≤ 0.05; *N*=4). C. Representative images showing the experimental setup used to sample buffer placed at the leaf-to-leaf contact points of plants (left) and a bar graph showing quantification of H_2_O_2_ at the buffer placed at the leaf-to-leaf contact points using the Amplex Red assay. Significance was determined with Student’s *t* test, *p ≤ 0.05, *N*=4. Scale bar is 1 cm. Abbreviations: CAT, catalase; CT, control; EL, excess light; R, receiver; ROS, reactive oxygen species; T, transmitter.

## Discussion

Our work reveals that in response to EL stress, plants that live in a group and physically touch each other exchange electric and Ca^2+^/ROS signals with each other, and that the transfer of Ca^2+^/ROS signals between different plants is required for their acclimation to this stress (Figs. 1-3, 11). In our previous work, we showed that the transfer of electric signals within a plant (*37*), or between different plants (*9*), is required for systemic, or P-T-P, ROS signaling to occur, respectively. Thus, mutations in the *GLUTAMATE-LIKE RECEPTOR* channels *GLR3.3* and *GLR3.6 (glr3.3glr3.6*) that abolish the transfer of systemic electric signals from local to systemic leaves within a plant, also block the transfer of Ca^2+^/ROS signals from one leaf to another (*37*). In addition, the inclusion of the *glr3.3glr3.6* double mutant as a mediator plant (using the same setup shown in Fig. 2), blocked the transfer of the P-T-P ROS signal from transmitter to receiver plants (*9*). It is therefore possible that the transfer of the electric signal from a transmitting plant to a receiving plant is required for the activation of the Ca^2+^/ROS signal in the receiver plant and that, in turn, the activation of the Ca^2+^/ROS signal in the receiving plant is required for acclimation (Fig. 11). Such a hierarchy in systemic signals was previously proposed to occur within a plant (*35*). The electric signal that rapidly spreads throughout the plant (*37*), and in the current study (Fig. 2), as well as in (*9*), also between different plants, could therefore cause depolarization of the plasma membrane, that excites and activates different calcium channels, causing an increase in cytosolic Ca^2+^ levels, that trigger ROS production (Fig. 11). In support of this model is our observation that the electric signal is very rapid and occurs within the first 0-2 min of EL stress application to the local leaf of the transmitter plant (even before the imaging process can be initiated; Fig. 2), and that the Ca^2+^/ROS signal appears in all plants at about the same time (and not spread from plant to plant in a successive manner; Fig. 2).

**Fig. 11.**
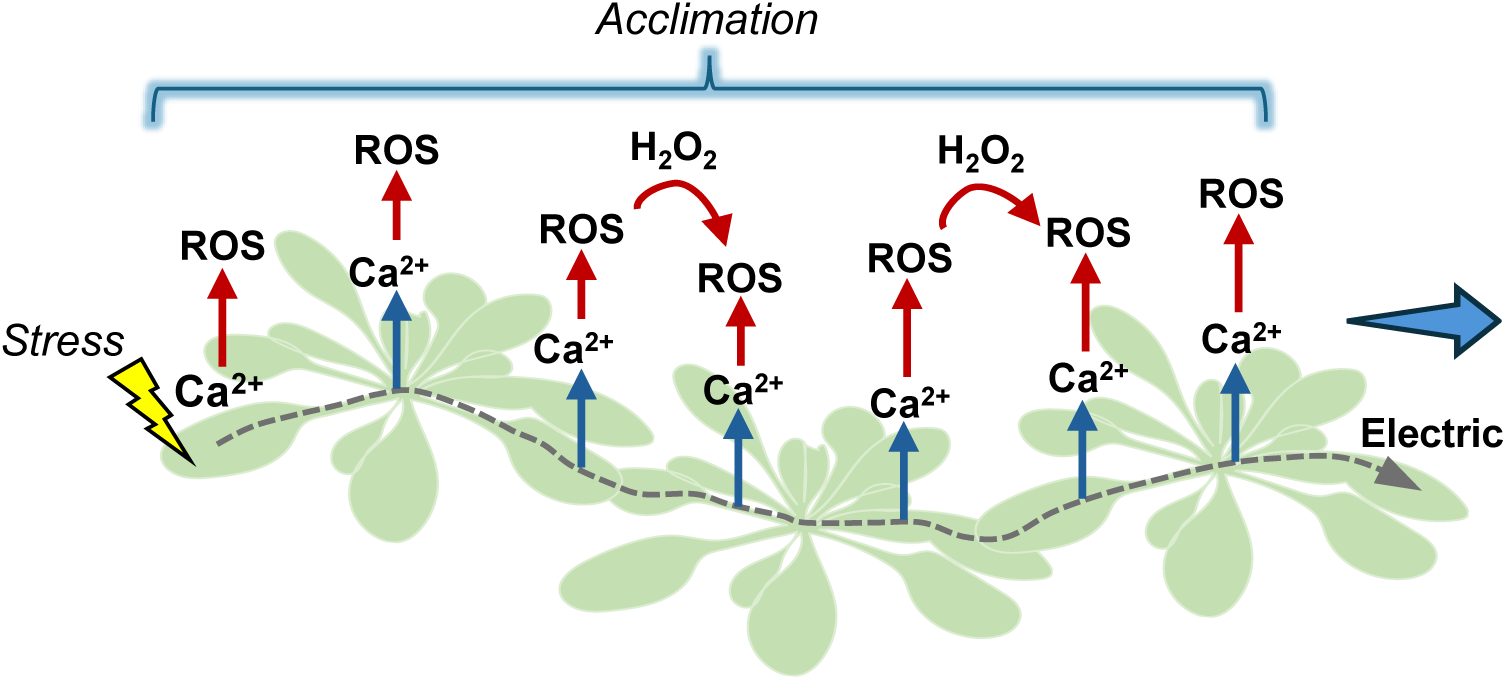
Proposed model for plant-to-plant signaling in response to stress. A localized stress (indicated with a jagged arrow) impacting a single leaf of a transmitting/initiating plant (on left) is shown to trigger the plant-to-plant (P-T-P) electric signal (dashed arrow) that is mobilized between plants and triggers Ca2+ (blue arrow) and ROS (red arrow) accumulation in each of the plants it travels through. Accumulation of H_2_O_2_ at the leaf-to-leaf contact points between plants is shown to be required for the successful mobilization of the P-T-P ROS signal between plants. All plants that have high Ca2+ and ROS levels are shown to undergo acclimation to the stress. A thick blue arrow (right) represents the direction of the P-T-P signal. Abbreviations: ROS, reactive oxygen species.

While the mechanism and hierarchy of P-T-P signaling described above may explain how the electric signal coordinates the triggering of the Ca^2+^/ROS signal among all plants physically touching each other (Fig. 11), it does not explain our findings with the *hpca1* mutant (Fig. 2). Thus, when the *hpca1* mutant is used as a mediator plant (*i.e.,* WT-*hpca1*-WT), the electric signal reached the receiver plant, but it did not activate the Ca^2+^/ROS signal in the receiver plant, as well as did not result in the enhanced acclimation of the receiver plant to EL stress (Figs. 2, 3). This finding suggests that the Ca^2+^ and/or ROS signals might also be directly mobilized between plants (a process that was blocked by the *hpca1* mutant that only transferred the electric signal; Fig. 2). While it might be harder to explain how calcium can be mobilized between two different plants touching each other (due to the constitutively high levels of calcium at the apoplast), H_2_O_2_, that can accumulate in the apoplast to very high levels (*38*, *39*), is known to easily diffuse in and out of cells (*35*, *36*, *40*, *41*). It is therefore possible that H_2_O_2_ that accumulates to high levels at a transmitting leaf of a high ROS producing transmitter plant will diffuse out of this leaf and into a receiving leaf of the low ROS producing receiver plant through their leaf-to-leaf contact point (as long as the electric wave was transferred between plants, and humidity is high enough to allow such a transfer). In support of this possibility are our findings that the presence of the H_2_O_2_ scavenger enzyme catalase at the leaf-to-leaf touching points suppresses the transfer of the ROS signal from plant to plant, and that H_2_O_2_ accumulates in an EL stress-dependent manner in the leaf-to-leaf contact points between plants (Fig. 10). While the transfer of H_2_O_2_ between different microorganisms of the same or different species is known to occur as part of different signaling and interaction mechanisms (*35*), limited evidence exists for the transfer of H_2_O_2_ between two different plants touching each other. More studies are therefore needed to address this intriguing possibility that may reveal a new mode of P-T-P signaling.

Our findings further suggest that plants that touch each other maintain higher levels of ROS/H_2_O_2_ compared to plants that live alone without a P-T-P physical touch (Fig. 9). Higher basal levels of H_2_O_2_ were also found in tomato (*Solanum lycopersicum*) plants inoculated with the plant growth-promoting rhizobacteria *Pseudomonas putida* (*34*). Interestingly, in both the tomato-*Pseudomonas* interaction (*34*) and the *A. thaliana* plants touching each other (Fig. 9) systems, application of stress to plants that already displayed higher basal levels of H_2_O_2_ resulted in the accumulation of even higher levels of H_2_O_2_. These findings could suggest that accumulating a higher basal level of H_2_O_2_ is part of the reason why plants that touch each other are more primed to acclimate to EL stress (Figs. 1, 9) and display higher steady-state levels of multiple stress response transcripts (Fig. 1). These findings are also in agreement with multiple studies that showed the potential benefits of higher basal H_2_O_2_ levels to cells, supporting proliferation and stress resilience in animals and plants (as long as these levels are not high enough to induce oxidative stress; *42*). The enhanced benefits of higher basal H_2_O_2_ levels to cells were termed by Dr. Helmut Sies as ‘oxidative eustress’ (*43*).

Although in our experimental setup the roots of plants that touched each other via leaf-to-leaf contacts did not touch each other (Figs. 1-8, 10; SI Appendix, Fig. S2; except for Fig. 9), we cannot rule out the possibility that volatiles and/or acoustic signals were also exchanged between plants. Further studies are needed to uncover the role of these communication pathways in inducing resilience to EL stress in a plant community. In this respect it should be noted that two independent alleles of a trichomeless mutant [*glabra1*; trichomes are major producers and storage sites for volatiles; (*44*, *45*)] that touched each other via leaf-to-leaf connections were able to exchange systemic ROS signals between each other (SI Appendix, Fig. S10), suggesting that the exchange of aboveground systemic signals reported here might not involve volatiles.

Taken together, our findings suggest that, compared to plants that do not physically touch or communicate with each other, plants growing in a dense community physically touching each other, in nature could exchange aboveground P-T-P signals that allow them to become more resilient to stress (Fig. 11). This advantage would of course come at the cost of less resources for growth, due to competition for water, nutrients, light, and/or space (*46*, *47*), but may offset such cost when the entire community experiences harsh environmental conditions (*48*). A tradeoff between acclimation stemming from living together and overall plant growth and reproduction may also occur within different agricultural settings, such as crop fields or plantations, in which crowding of plants could result in yield penalty (*49*, *50*). Thinning a dense forest population of Norway spruce saplings was recently shown, for example, to result in a long-term ‘high physiological stress’ state among the surviving saplings within the thinned community (*51*), supporting our findings. As environmental conditions on our planet are worsening due to climate change and increased pollution levels (*52*, *53*), that generate conditions of multifactorial stress combination (*54*, *55*), our findings may offer a promise for future agriculture and ecosystem engineering, highlighting the potential of using plant communities with enhanced P-T-P signaling for heightened resilience (*56*). In this respect, it should be noted that a key acclimation mechanism of the unicellular green algae *Chlamydomonas reinhardtii* to conditions of multifactorial stress combination, identified recently, is the formation of cell aggregates (*57*, *58*). At least one possible answer to the growing complexity of environmental stress conditions on our planet could therefore be living together. In addition to humans and some animals, the principle of ‘United We Stand’ may hence also apply to plants living on Earth.

## Materials and Methods

### Plant material, growth conditions, and stress treatments

*Arabidopsis thaliana* wild-type Col-0 (WT) and mutant (Dataset S11) plants were germinated and grown on peat pellets (Jiffy 7; Jiffy International) under controlled conditions of 10-hour/14-hour light/dark regime, 50 μmol photons s^−1^ m^−2^, and 21°C for 4 to 5 weeks. Leaf-to-leaf connections between different plants were established by allowing plants to physically touch each other in the absence of any agar or buffer solution, however, following treatment of plants with the imaging buffer (see below; *9*). Leaf-to-leaf connections between different plants were also established by adding a drop of a solution containing 0.05 M phosphate buffer, pH 7.4, 0.01% (v/v) Silwet L-77 and 0.05% (w/v) agarose, containing the different dyes used for imaging (*9*). No differences were found in P-T-P ROS signaling between the different setups, including when Silwet L-77 and agar were omitted from buffers (SI Appendix, Fig. S11). In almost all the setups we used, only one leaf from each plant touched a similar shape and size leaf from a different plant. Connections were maintained for different periods of time depending on the specific experiment. To induce a local EL stress treatment in individual plants, or plants connected to one another in a community, a single leaf (local, L) of the transmitter plant was illuminated with 2000 μmol photons s^−1^ m^−2^ using a ColdVision fiber optic LED light source (Schott) for 10 min (SI Appendix, Fig. S2). For each experiment, plants of the same age and developmental stage were used, and all imaging or sampling were conducted between 9AM to 1PM.

### RNA-Seq analysis

RNA-seq analysis, comparing individual plants to a group of plants that touch each other aboveground, followed the experiment designed shown in Fig. 1A (similar to control plants shown in SI Appendix, Fig. S1 but in triplets). WT plants were grown as described for 4 weeks. Plants were then divided into three groups: (i) For individual plants, triplets of individual plants were maintained at a distance of 2.5 cm from each other (measured from leaf edges) with no leaves connected; (ii) and (iii), for communities, two leaves of each plant were connected with two other plants (forming a triangle) as explained above (Fig. 1A; SI Appendix, Fig. S1). Triplets of connected plants as well as triplets of individual plants were maintained under control conditions for 1 hour (ii) or 1 week (iii). Three non-connecting leaves per plant were sampled and flash frozen in liquid nitrogen in three replicates and a total of 10 plant triplets per condition were collected. All plants used for the experiments were of the same age and developmental stage and sampled on the same day (between 10-11AM). For the RNA-seq analysis involving excess light (EL) stress treatment applied to a single leaf (local leaf; L), WT and *hpca1* plants were grown as described above for 4 weeks. Plants were then organized in two different setups as shown in Fig. 3A (SI Appendix, Fig. S2), allowing a single leaf from each plant to touch the other: (i) a chain of three WT (Col-0) plants touching each other consisting of a transmitter (T), mediator (M), and receiver (R) plants (WT-WT-WT), and (ii) a chain of three plants (T, M, R) in which the T and R plants were WT (Col-0), and the M plant an *hpca1* mutant (WT-*hpca1*-WT). Leaves of plants were connected as described above for 0.5 hour before the EL stress treatment. One leaf (local, L) of the T plant was subjected to excess light stress (1700 μmol photons s^-1^ m^-1^ for 30 min). Local and systemic leaves (L, S1, S3; Fig. 3A) from control and treated plants were sampled. Leaves from 60 different plants were pooled into three replicates, and every replica included 20 leaves/plants for each sample (L, S1, S2 and S3). Samples from the two experiments described above were processed similarly. RNA isolation, sequencing, quality control, read trimming, alignment, differential transcript expression, and principal component analysis were conducted according to (*21, 22, 59*–*63*). RNA-Seq data files are available at GEO (GSE301070). Transcript datasets of stress, hormone, and ROS responses used in Figs. 1D and 3D were obtained from (*64*) with some updates as follows: transcripts associated with a ‘touch’ response were obtained from (*65*); transcripts related to ‘excess light’ (EL) response were obtained from (*22*) and transcripts from ‘waterlogging’ list were filter by fold change ≤ -1 and ≥ 1 (*64*). Venn diagrams were created in VENNY 2.1 (BioinfoGP). UpSet plots were generated using the UpSetR (*66*) package.

### GRN analysis

Gene regulatory network inference analysis was conducted using the DIANE computational framework, which integrates normalization, differential expression analysis, expression-based clustering, and machine-learning-driven network reconstruction into a unified analytical pipeline (*67*). The raw count matrices from leaf S3 of each setup were imported into DIANE and processed under the same experimental design structure described above to ensure that the datasets are compared to one another. Normalization by Dseq2 was used to remove low-abundance transcripts, followed by quality control steps, which confirmed the expected separation of the transcriptomic signatures between setups. Differential expression analysis was carried out with the edgeR engine integrated into DIANE. Negative binomial dispersion estimates were prepared for each dataset, while generalized linear models were fitted to assess transcriptional changes associated with EL treatment in leaf S3. A separate list of DEGs was generated for the WT-WT-WT S3 samples and the WT-*hpca1*-WT S3 samples. Functional enrichment was conducted using ClusterProfiler in DIANE, which allowed the identification of biological processes and molecular pathways that underpin EL-induced transcriptional regulation in each configuration. Expression-based clustering was conducted using DIANE’s Coseq mixture model algorithm. Clusters were divided based on dynamic expression patterns in each data set, which resulted in statistically coherent expression modules for EL-responsive signaling in S3. These modules formed the structural basis of network inference. GRN reconstruction was performed based on the implementation of different algorithms in DIANE. For each setup, gene expression was modeled through random forest regression, where the expression of a target gene is predicted by the expression profiles of all remaining genes. Feature-importance measures were computed to infer high-confidence regulatory edges, and DIANE retained the top-ranked connections to generate condition-specific S3 subnetworks. All DIANE-generated subnetworks were imported into Cytoscape version 3.12 for advanced visualization and layout refinement.

### Anthocyanin content quantification

Whole individual plants or communities composed of four plants (touching each other for 1 hour) were exposed to excess light conditions (2000 μmol photons s^−1^ m^−2^) using a full spectrum led grow light (BESTVA, CA, USA) positioned 22 cm above plants as shown in SI Appendix, Fig. S1 for 24 hours. Control plants were grown in parallel and sampled at the same time. Whole rosettes were collected independently (flash frozen in liquid nitrogen), and each plant was treated as a technical replicate. Anthocyanin content was determined as described in (*68*). Leaf temperature (20-22°C) was monitored as described in (*21*).

### H_2_O_2_ measurements

H_2_O_2_ quantification in individual plants and in large plant communities was performed using Amplex-Red (10-acetyl-3,7-dihydroxyphenoxazine [ADHP]; Thermo Fisher Scientific, Waltham, MA, USA) as described by (*18*). Col-0 seeds (5 for individual and ˃100 for communities), were sown on 8.5 cm x 12.5 cm pots filled with Promix BX growing medium (Premier Tech Horticulture, Quakertown, PA, USA). Plants were grown under controlled conditions, as previously described, for 3 to 4 weeks. Leaves and roots were permitted to grow without spatial restriction. Leaves from individual plants and plants living in communities were collected and immediately frozen and ground to fine powder, resuspended in 100 μL 0.1 M TCA (Thermo Fisher Scientific, Waltham, MA, USA), and centrifuged for 15 min at 12,000 × *g*, 4 °C. The supernatant was buffered with 1 M phosphate buffer pH 7.4, and the pellet was dried and used for dry weight calculation. H_2_O_2_ quantification in the supernatant was performed according to the MyQubit-Amplex-Red Peroxide Assay manual (Thermo Fisher Scientific, Waltham, MA, USA), using a calibration curve of H_2_O_2_ (Thermo Fisher Scientific, Waltham, MA, USA) as described by (*18*). Four leaves from five different plants were pooled together in one replicate and a total of five replicates were processed. For H_2_O_2_ quantification at the leaf-to-leaf contact point between two different plants, plants were fumigated with a solution containing 0.05 M phosphate buffer, pH 7.4, 0.01% (v/v) Silwet L-77 for 30 min. Leaves were then connected with 200 μL of the same solution plus 0.05% (w/v) agarose using a microscope cover glass as support. To induce a local EL stress treatment, a single leaf of the transmitter plant was illuminated with 2000 μmol photons s^−1^ m^−2^ using a ColdVision fiber optic LED light source (Schott) for 10 min. After EL stress, control and treated plants were maintained under the same conditions for 30 minutes before sampling. 100 μL of the solution at the leaf-to-leaf contact point was recovered and used for H_2_O_2_ quantification. Samples were mixed with 100 μL 0.1 M TCA (Thermo Fisher Scientific, Waltham, MA, USA), and centrifuged for 15 min at 12,000 × *g*, 4 °C. The supernatant was buffered with 1 M phosphate buffer pH 7.4 and H_2_O_2_ quantification was performed as previously explained. The reaction was conducted in dark for 45 min-1 hour instead of 30 min. A calibration curve of H_2_O_2_ prepared in the solution applied to the leaf-to-leaf contact point between different plants was included. Four pairs of plants per treatment and three independent experiments were processed.

### Acclimation assays

Acclimation was measured in individual plants as well as in plants that touched each other connected by one leaf (SI Appendix, Fig. S4). For individual plants, systemic acclimation to excess light stress was tested by exposing a local leaf to excess light (2000 μmol photons s^−1^ m^−2^) for 10 min, incubating the plant under controlled conditions for 50 min, and then exposing the same leaf (local) or another younger leaf (systemic) to excess stress (2000 μmol photons s^−1^ m^−2^) for 45 min (SI Appendix, Fig. S4B), as previously described (*18*, *21*, *22*). For measuring acclimation in a chain of plants, as shown in Fig. S4A, plants were misted with water for 10 min and leaves were connected as described above for 30-60 min. Chains of plants composed of WT(T)-WT(M)-WT(R), as well as chains of plants composed of WT(T)-*hpca1*(M)-WT(R) were treated side by side. For non-acclimated chains, excess light was applied to a systemic leaf (S3) of the receiver plant (WT in both setups) for 45 min with no stress pre-treatment (*i.e.,* no EL-pretreatment was applied). For acclimated chain of plants, a short pre-treatment (10 min) of excess light (2000 μmol photons s^−1^ m^−2^) was applied to a local (L) leaf of the transmitter plant (WT in both setups). Following the pre-treatment, plants were allowed to rest for 2 hours to induce acclimation. Excess light (2000 μmol photons s^−1^ m^−2^) was then applied to a systemic leaf (S3) of the receiver plant (WT) for 45 min. Control plants were maintained under the same control conditions and sampled at the same time. For the three groups, the local (L) leaf and a systemic leaf (S1) of the transmitter (T) plant, together with the systemic (S3) leaf of the receiver (R) plant were sampled for conductivity measurement (described below). Leaf injury after excess light stress was measured using the electrolyte leakage assay, as described previously (*18*, *21*, *22*). For whole plants exposed to excess light, either individually or in a community, stress was applied for 24 hours following the setup shown in SI Appendix, Fig. S1 as explained above. Following excess light treatment, one non-connected leaf per plant was sampled and placed in a tube containing 30 mL of ddH_2_0 and moved to a gentle shaker for one hour. After one hour, the electrolyte leakage was measured (*18*, *21*, *22*).

### Whole-plant imaging of ROS, calcium and membrane potential

As previously described (*18, 37*), plants were treated with fluorescent dyes via fumigation (misting with fine droplets) in a glass container using a nebulizer (Punasi Direct, Hong Kong, China) for 30 minutes with: 50 µM H_2_DCFDA (Millipore-Sigma, St. Louis, MO, USA) for ROS imaging; 4.5 µM Fluo-4-AM (Becton, Dickinson and Company, Franklin Lakes, NJ, USA) for calcium imaging; 20 µM DiBAC_4_(3) (Biotium, Fermont, CA, USA) for imaging changes in membrane potential. Following fumigation, a chain of physically connected plants was established, consisting of a transmitter (T) and a receiver (R) WT plants that were separated by a mediator plant (M), that was either WT, *hpca1,* or *gl1* mutant (Dataset S11). Transmitter plants were then treated with a local excess light (EL; 2000 μmol photons s^−1^ m^−2^) for 10 min and imaged using the IVIS Lumina S5 fluorescence imaging system (PerkinElmer, Waltham, MA, USA) as previously described (*18, 37*). Fluorescence images (excitation/emission 480/520 nm) were acquired every minute for 30 min. Accumulation of ROS, calcium and electric (membrane potential) signals, in treated and untreated plants, were compared to the 0-min time point and determined by subtracting the fluorescent signal of the initial time point (0-min) from the time point of interest. The 0-min time point was the initial image, which served as a baseline, resulting in no visible signal after subtraction using the Living Image 4.7.2 software (PerkinElmer). The total radiant efficiency of regions of interest (ROI; indicated with white circles in the different Figures) of only non-plant-connected leaves were quantified as pixel intensity (Raw Int Den) using ImageJ. Each experiment was repeated at least three times, *N* for each experiment is indicated in the Figure legends. For imaging ROS accumulation in large plant communities, Col-0 seeds (5 for individual and ˃100 for communities), were sown on 8.5 cm x 12.5 cm pots filled with Promix BX growing medium (Premier Tech Horticulture, Quakertown, PA, USA). Plants were grown under controlled conditions, as previously described, for 3 to 4 weeks. Leaves and roots were permitted to grow without spatial restriction. Plant communities were fumigated with the fluorescent dye H_2_DCFDA for 30 minutes, as described above. After fumigation, plants were maintained under the same controlled conditions for an additional hour before stress application. Excess light stress (2000 μmol photons s^−1^ m^−2^) was applied to a local region (indicated with a yellow circle in Fig. 9C) for 10 min and plant communities were imaged using the IVIS Lumina S5 fluorescence imaging system. Fluorescence images (excitation/emission 480/520 nm) were acquired every minute for 30 min. Image processing was conducted as described above, and two distinct regions of interest (S1 and S2, marked with white rectangles in Fig. 9) were quantified using ImageJ. Fluorescence intensity was measured as pixel intensity (Raw Int Den). To study the role of H_2_O_2_ in P-T-P signaling, the H_2_O_2_ scavenger enzyme catalase (2000 U/ml final concentration; Millipore-Sigma) was added to the phosphate buffer (described above) at the leaf-to-leaf contact point between two different plants. Dye fumigation, excess light treatment and imaging of accumulation of ROS in plants with leaves connected with the solution, containing or not, the scavenger catalase were performed as described above.

### Statistics and data analysis

Data analysis and statistics were performed in GraphPad Prism 10. Different letters indicate significant differences (p ≤ 0.05) as determined by a one-way ANOVA followed by uncorrected Fisher’s LSD post hoc tests, and results are presented as means ± standard errors (SE). Asterisks denote statistically significant differences based on two-sided Student’s *t* test with significance levels: *p ≤ 0.05; **p ≤ 0.01; ***p ≤ 0.001. Asterisks in Fig. 9C indicate significant differences (p ≤ 0.05) as determined by a two-way ANOVA followed by Tukey’s HSD post hoc test. Differentially expressed transcripts were defined as those that had a fold-change with an adjusted p ≤ 0.05 (negative binomial Wald test followed by Benjamini–Hochberg correction).

## Acknowledgments

We thank the Arabidopsis Biological Resource Center (https://abrc.osu.edu/) for seeds of Arabidopsis mutants used in this study.

## Funding

National Science Foundation, IOS-2414183 (RM).

National Science Foundation, IOS-2110017 (RM).

National Science Foundation, IOS-2343815 (RM, TJ).

Polish National Science Center, OPUS20, UMO-2020/39/B/NZ3/02103 (SMK).

## Author contributions

Conceptualization: RM, MAPV, YF, SMK.

Methodology: MAPV, YF, AG, SD, AB, RA.

Investigation: MAPV, YF, AG, SD, RM.

Visualization: MAPV, YF.

Funding acquisition: RM, TJ, SMK, RA.

Project administration: RM.

Supervision: RM, TJ, RA.

Writing – original draft: RM, MAPV.

Writing – review & editing: RM, MAPV, YF, AG, SD, SMK, TJ.

## Data and materials availability

All materials used in the analysis are available upon request from the Corresponding Author. All data are available in the main text or the supplementary materials. RNA-Seq data is available at Gene Expression Omnibus (GEO) under accession GSE301070. Supplementary Datasets for this study were deposited in https://datadryad.org/ under DOI: 10.5061/dryad.jsxksn0q9.

## Supplementary Figures

**Fig. S1.**
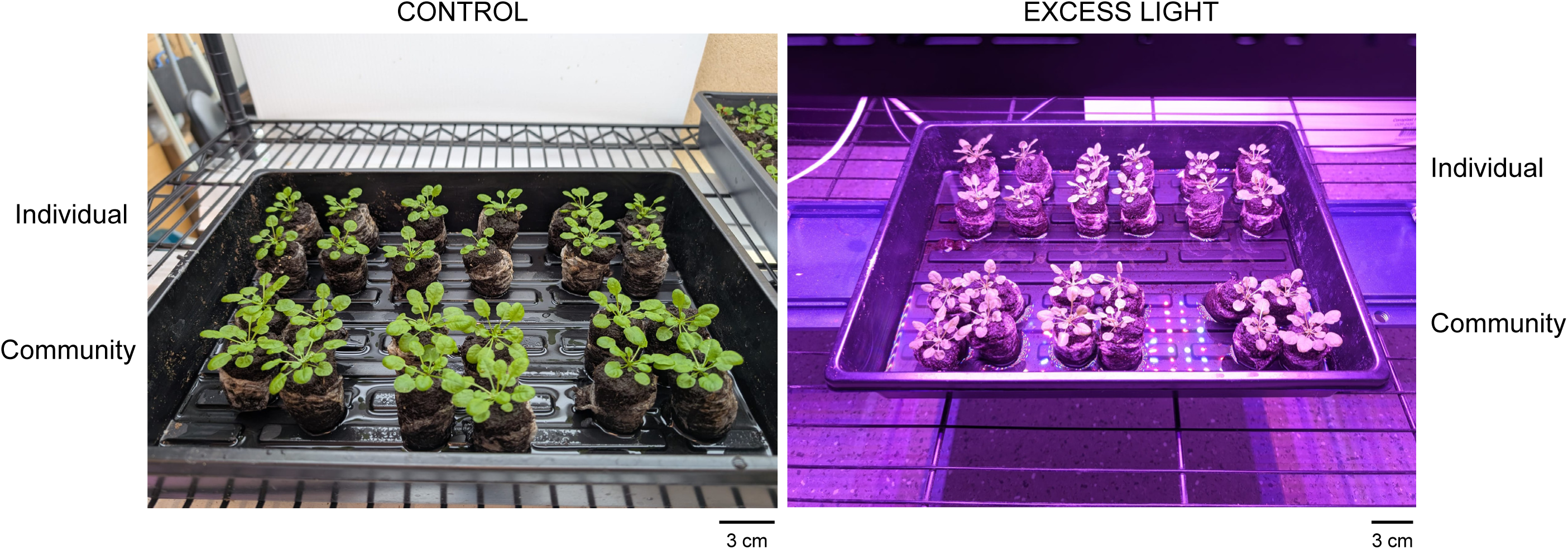
Images showing the experimental setup used to compare the resilience of individual *A. thaliana* plants to that of groups of *A. thaliana* plants that physically touch each other aboveground, to excess light stress. Individual plants were maintained at a distance of 2.5 cm from each other and groups of plants (touching each other) were connected for 1 hour by a drop of a solution containing water, or 0.05 M phosphate buffer, pH 7.4, 0.01% (v/v) Silwet L-77 and 0.05% (w/v) agarose, to ensure continuous contact between touching leaves. Quadruplets of individual plants were maintained under control or excess light stress conditions for 24 hours and then sampled for quantification of anthocyanin content and leaf injury. In support of Fig. 1.

**Fig. S2.**
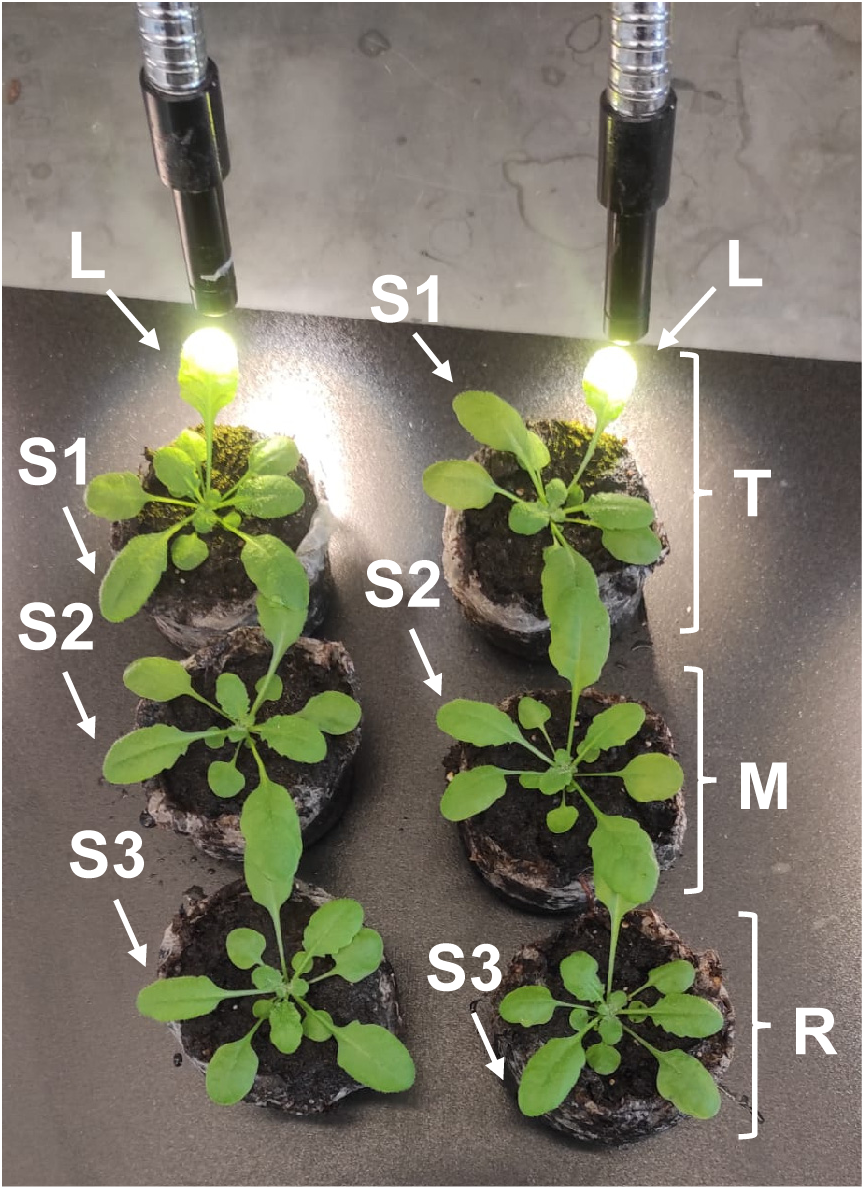
Image showing the experimental setup used to study the transfer of plant-to-plant signals (electric, calcium and reactive oxygen species) between three successively arranged plants touching each other, in response to excess light (EL) stress applied to one leaf of the transmitter plant. Leaf-to-leaf connections between different plants were established by adding a drop of water or a solution containing 0.05 M phosphate buffer, pH 7.4, 0.01% (v/v) Silwet L-77 and 0.05% (w/v) agarose, containing the different dyes used for imaging, to ensure continuous contact between touching leaves. Following 30-60 min incubation, EL stress was applied to one leaf of the transmitter plant. Transmitter (T), mediator (M), and receiver (R). In support of Fig. 2.

**Fig. S3.**
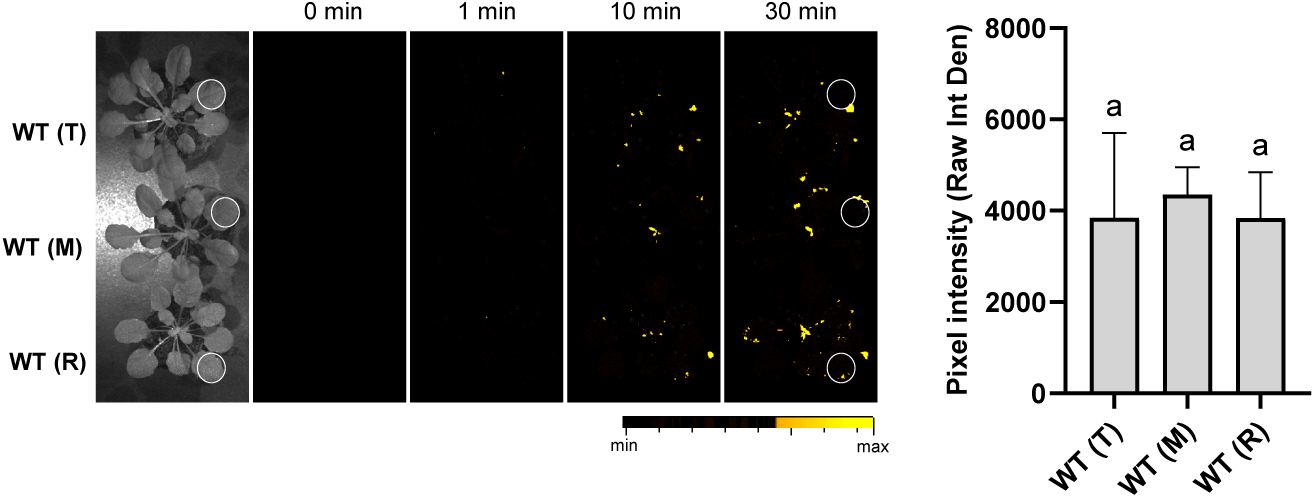
The plant-to-plant (P-T-P) reactive oxygen species (ROS) wave is not transmitted between plants under non-stress conditions. A representative image showing the experimental setups used to study the transfer of the P-T-P ROS signal between three successively arranged plants touching each other (transmitter – T, mediator – M, and receiver – R) and time lapse images of ROS accumulation in plants touching each in the absence of an excess light (EL) stress treatment are shown on left, and a bar graph showing statistical analysis of ROS accumulation at 30 min (*N*=3) is shown on right. Significance was determined by one-way ANOVA followed by a Fisher’s LSD post hoc test (different letters denote statistical significance at p ≤ 0.05). In support of Fig. 2. Control (CT), Transmitter (T), mediator (M), and receiver (R).

**Fig. S4.**
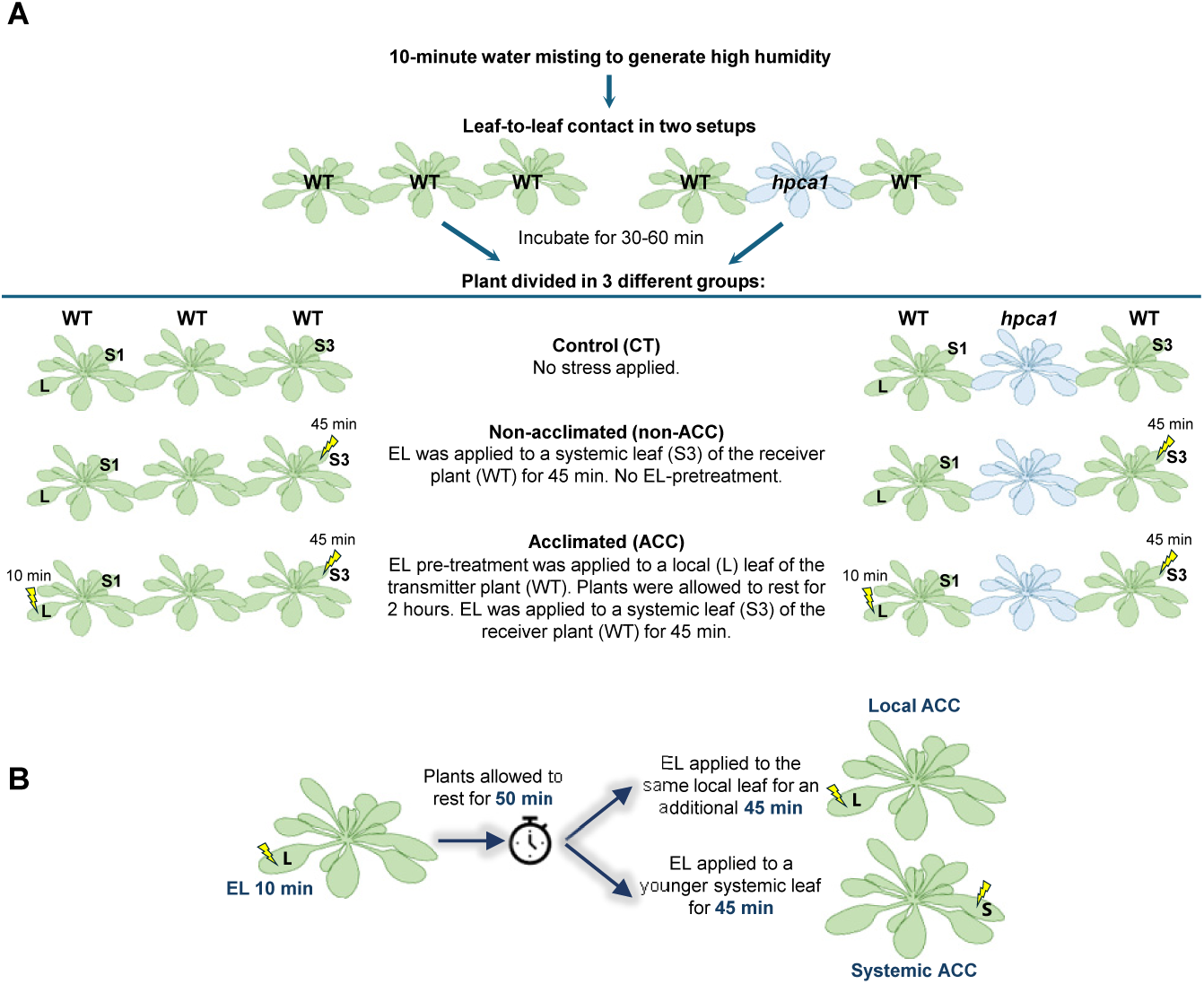
Experimental setup used to study local (L) and systemic (S) leaf acclimation to excess light (EL) stress in three successively arranged plants touching each other (A) or in individual plants (B). In support of Figs. 3 and 6.

**Fig. S5.**
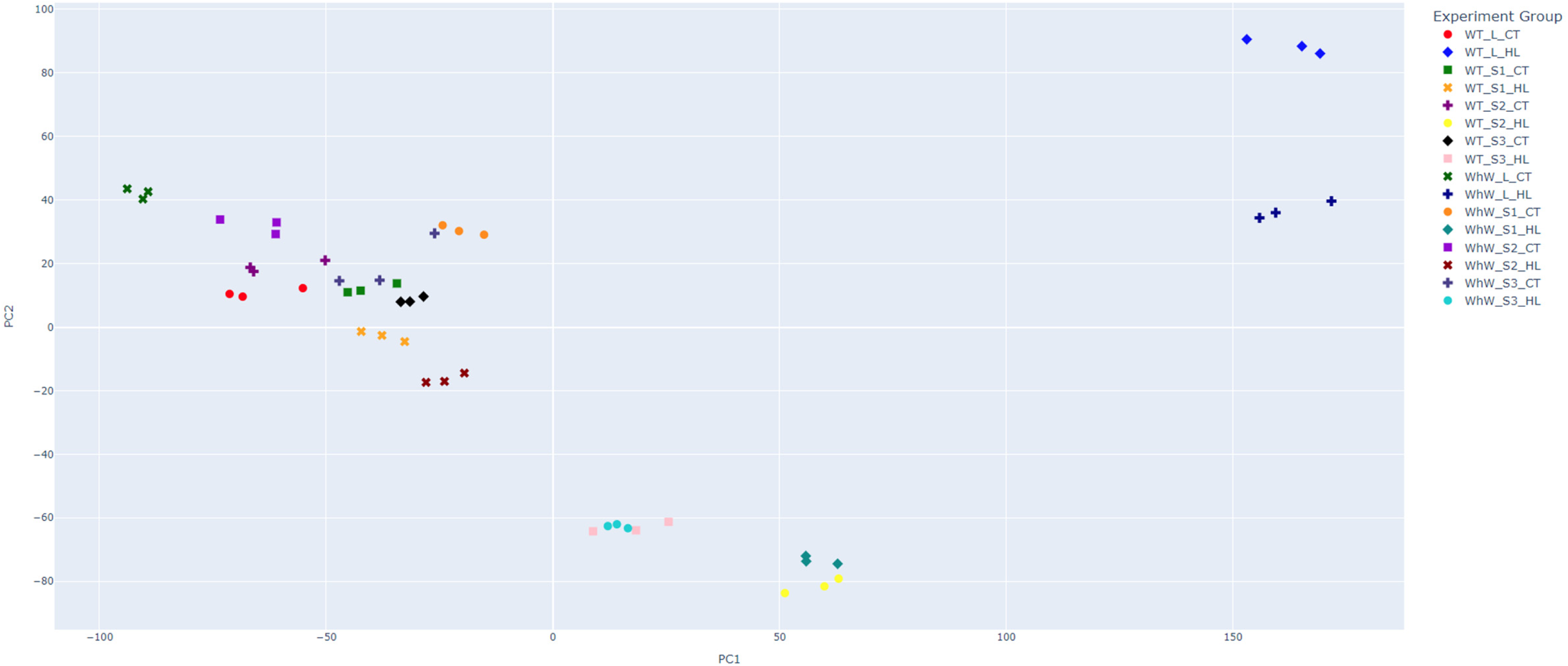
A PCA plot showing the distinct transcriptomics grouping of plants arranged in the WT-WT-WT and WT-*hpca1*-WT setups in response to excess light (EL) stress applied to a single leaf of the transmitter plant. In support of Fig. 4. Abbreviations: WT, WT-WT-WT setup; WhW, WT-*hpca1*-WT setup; L, local leaf; S1, systemic leaf 1 of the transmitter plant; S2, systemic leaf 2 of the mediator plant; S3, systemic leaf 3 of the receiver plant; CT, control; HL, excess light (EL) stress.

**Fig. S6.**
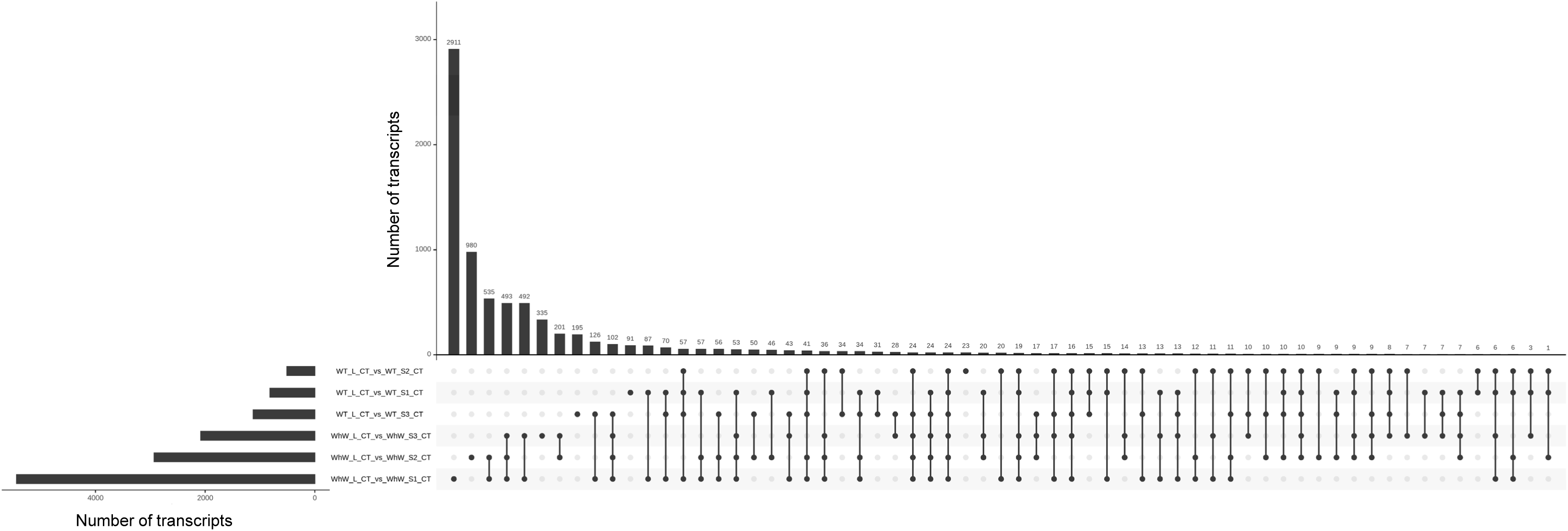
UpSet plot showing the unique and common transcripts significantly altered in their expression under control conditions (CT), in the systemic leaves 1 (S1), 2 (S2) and 3 (S3) compared to local leaf (L), in the WT-WT-WT (WT) and WT-*hpca*1-WT (WhW) setups. In support of Fig. 4.

**Fig. S7.**
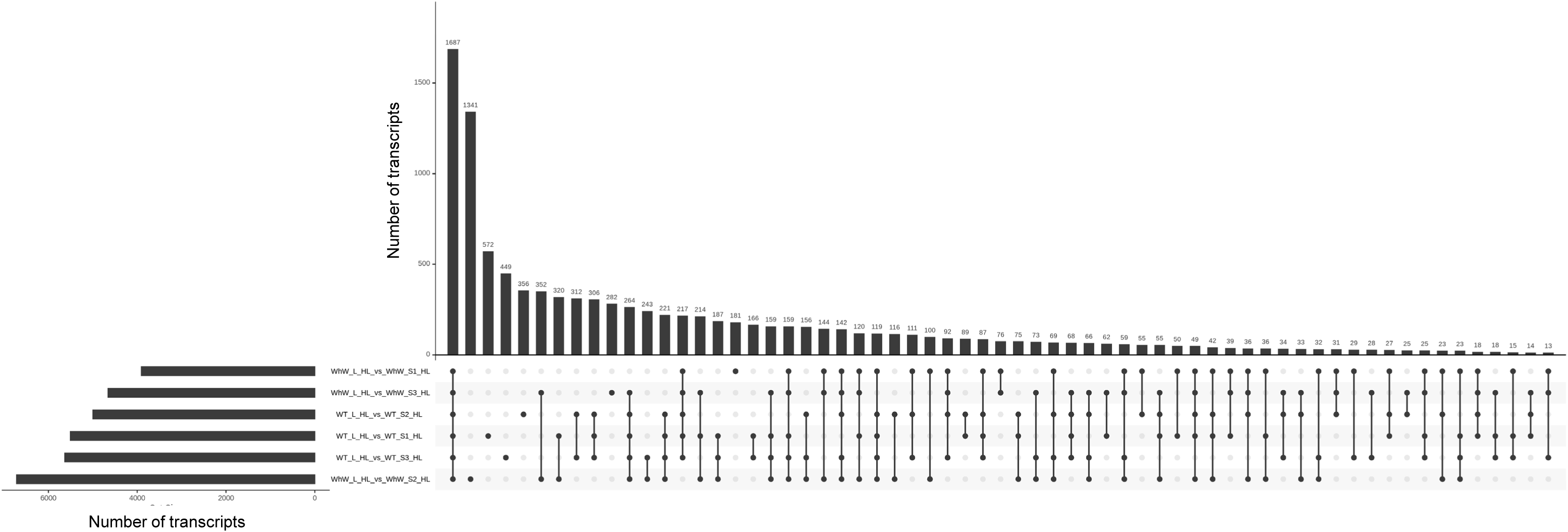
UpSet plot showing the unique and common transcripts significantly altered in their expression in the systemic leaves 1 (S1), 2 (S2) and 3 (S3) compared to local leaf (L) in the WT-WT-WT (WT) and WT-*hpca1*-WT (WhW) setups in response to excess light/high light (HL) stress applied to the local leaf of the transmitter plant. In support of Fig. 4.

**Fig. S8.**
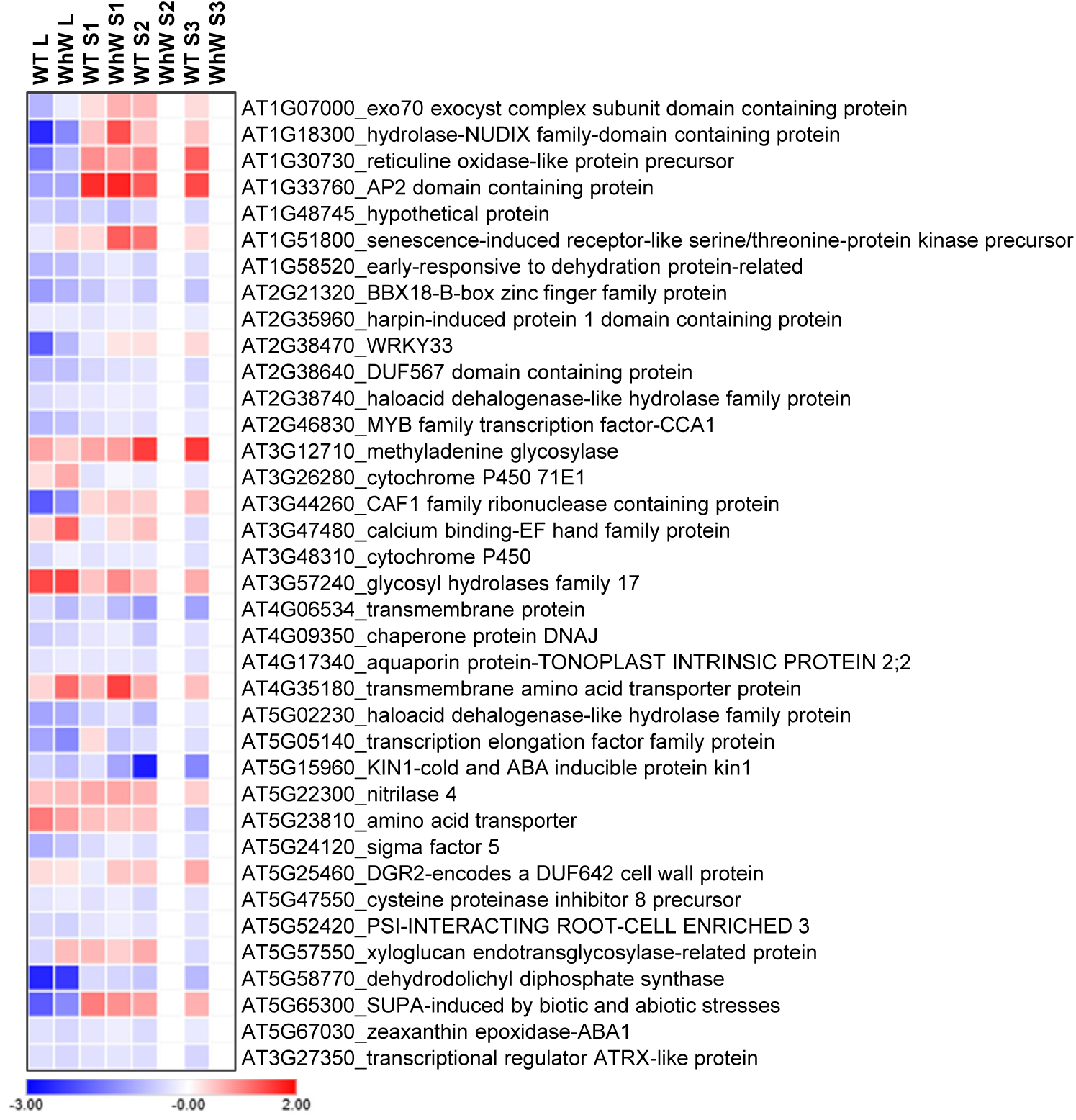
Heat map showing the expression pattern (fold change) of all putative calcium/reactive oxygen species-dependent transcripts significantly altered during plant-to-plant signaling in response to a local application of excess light (EL) stress applied to the local leaf of the transmitter plant. Abbreviations: WT, WT-WT-WT setup; WhW, WT-*hpca1*-WT setup; L, local leaf; S1, systemic leaf 1 of the transmitter plant; S2, systemic leaf 2 of the mediator plant; S3, systemic leaf 3 of the receiver plant. In support of Fig. 5.

**Fig. S9.**
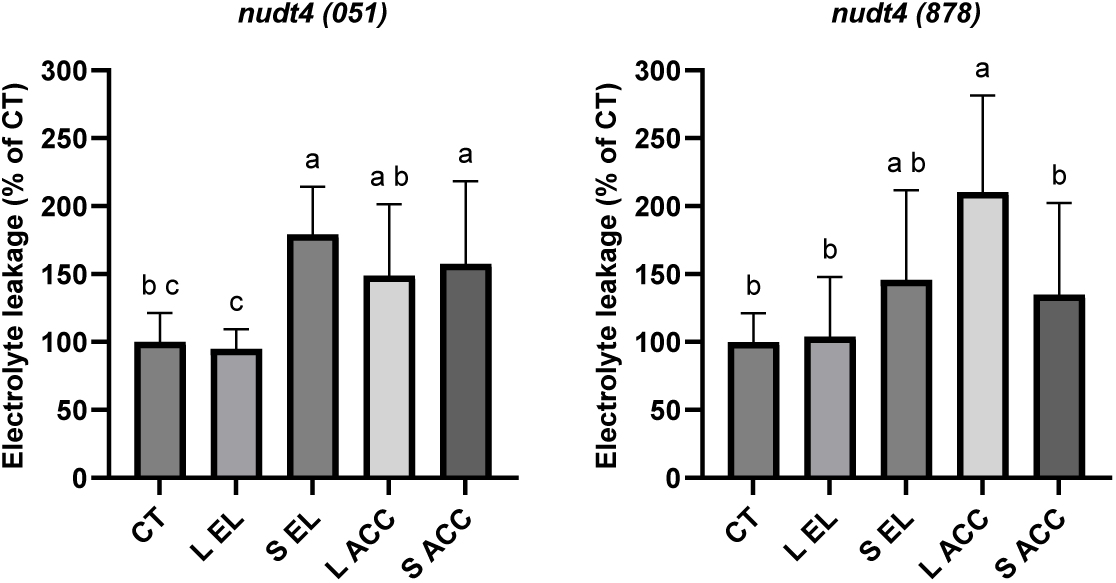
Cellular damage (measured as ion leakage) in local (L) and systemic (S) leaves of two knockout mutants of the nudix hydrolase 4 gene (*NUDT4; 69*) subjected to an extended excess light (EL) stress following a short (10 min) pretreatment of EL stress. Significance was determined by one-way ANOVA followed by a Fisher’s LSD post hoc test (different letters denote statistical significance; p ≤ 0.05; *N*=5). In support of Fig. 6. Abbreviations: acclimated (ACC), control (CT).

**Fig. S10.**
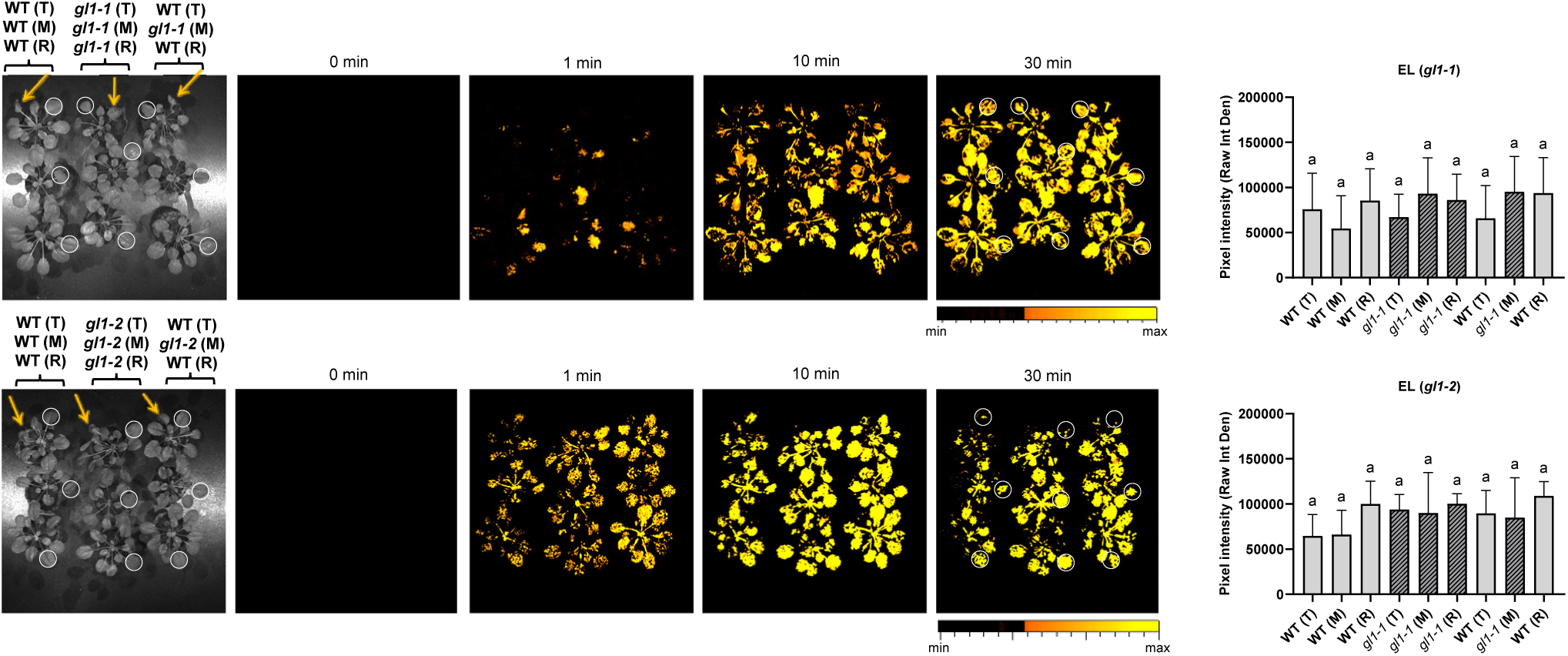
The systemic reactive oxygen species (ROS) plant-to-plant (P-T-P) signal can be transmitted between wild type and trichomeless mutants, or from one trichomeless mutant to another, in response to excess light (EL) applied to a local leaf of a transmitter plant (wild type or a trichomeless mutant). Images showing the experimental setups used to study the transfer of the P-T-P ROS signal between three successively arranged plants touching each other (transmitter – T, mediator – M, and receiver – R; Left), time lapse images of ROS accumulation in plants touching each in response to an EL stress treatment (0-30 min; Middle), and bar graphs showing statistical analysis of ROS accumulation at 30 min (Right; *N*=3). The different setups used included two independent alleles of a trichomeless mutant (*glabra1*; *gl1-1*; Top; and *gl1-2*; Bottom). White circles indicate regions of interest used to quantify signal intensity. Significance was determined by one-way ANOVA followed by a Fisher’s LSD post hoc test (different letters denote statistical significance at p ≤ 0.05). In support of Fig. 2.

**Fig. S11.**
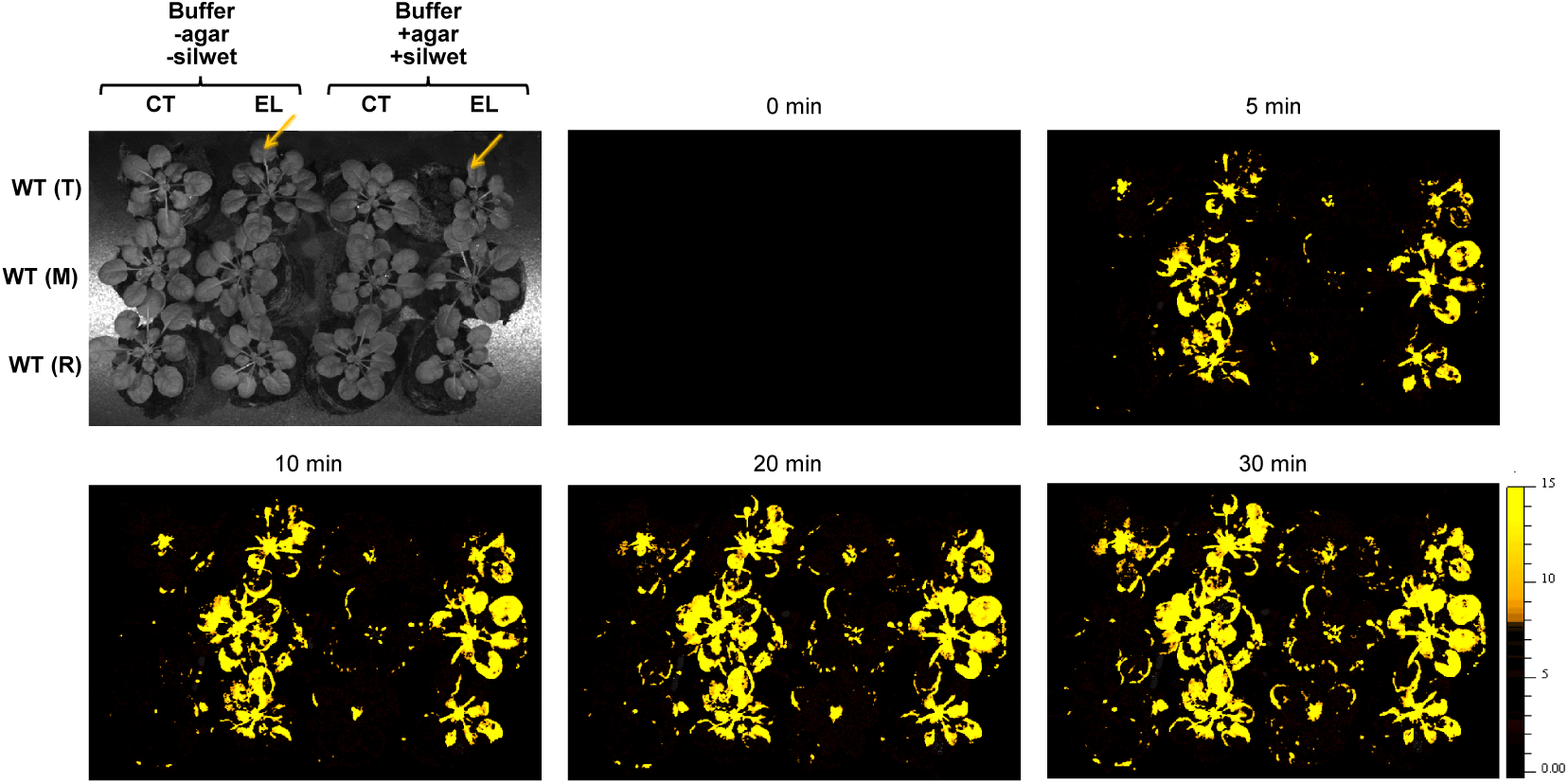
Images showing the experimental setups used to study the transfer of the plant-to-plant (P-T-P) reactive oxygen species (ROS) signal between three successively arranged plants touching each other (transmitter – T, mediator – M, and receiver – R). The two different setups used included plants connected with a drop of a solution containing 0.05 M phosphate buffer, pH 7.4 (Buffer – agar –silwet), or the same solution plus 0.01% (v/v) Silwet L-77 and 0.05% (w/v) agarose (Buffer +agar +silwet). Arrows indicate the leaf subjected to excess light stress. In support of Fig. 2. Abbreviations: CT, control; EL, excess light; M, mediator; R, receiver; T, transmitter.

**Datasets S1-S15** (Deposited in https://datadryad.org/ under DOI: <u>10.5061/dryad.jsxksn0q9</u>)

**Dataset S1.** Transcripts significantly altered in their expression in wild type *Arabidopsis thaliana* plants growing individually (individual) or in a community touching each other (community) for 1 hour or 1 week. In support of Fig. 1.

**Dataset S2.** Overlap between transcripts significantly altered in their expression in plants growing as a community and touching each other for 1 hour or 1 week, compared to plants growing individually. In support of Fig. 1C.

**Dataset S3.** Percent representation of transcripts related to different stress-, hormone-, and reactive oxygen species (ROS)-response (left column) among transcripts significantly altered in their expression in plants living as a community and touching each other, compared to individual plants. The number of transcripts is shown in parentheses. In support of data presented in Fig. 1C-D.

**Dataset S4.** Transcripts significantly altered in wild type and the mutant *hpca1* in response to excess light stress applied to a local leaf of the transmitter plant in the two setups WT-WT-WT and WT-*hpca1*-WT. WT-WT-WT setup is abbreviated as ‘WT’ and WT-*hpca1*-WT setup as ‘WhW’; ‘L’ is local leaf, ‘S1’ is systemic leaf 1 of the transmitter plant, ‘S2’ is systemic leaf 2 of the mediator plant, ‘S3’ is systemic leaf 3 of the receiver plant, ‘CT’ is control and ‘EL’ is excess light.

**Dataset S5.** Overlap between transcripts significantly altered in their expression in local (L) and systemic (S1-S3) leaves of plants from two setups (WT-WT-WT, abbreviated as ‘WT’; WT-*hpca1*-WT, abbreviated as ‘WhW’) in response to excess light (EL) stress applied to the local leaf of the transmitter plant. In support of Fig. 4B. Control, CT.

**Dataset S6.** Overlap between transcripts significantly altered in their expression in local (L) and systemic (S1-S3) leaves of plants from two setups (WT-WT-WT, abbreviated as ‘WT’; WT-*hpca1*-WT, abbreviated as ‘WhW’) in response to excess light (EL) stress applied to the local leaf of the transmitter plant. In support of Fig. 4C.

**Dataset S7.** Percent representation of transcripts related to different stress-, hormone-, and reactive oxygen species (ROS)-response (left column) among transcripts significantly altered in their expression in systemic leaf 1 (S1) from the transmitter plant in the setup WT-*hpca1*-WT in response to excess light stress applied to the local leaf of the transmitter plant. The number of transcripts is shown in parentheses. In support of data presented in Fig. 4.

**Dataset S8.** List of putative calcium/reactive oxygen species (ROS)-dependent transcripts significantly altered in their expression in plants during P-T-P communication. In support of data presented in Fig. 5.

**Dataset S9.** List of putative electric (membrane potential)-dependent transcripts significantly altered in their expression in plants during P-T-P communication. In support of data presented in Fig. 5.

**Dataset S10.** Percent representation of transcripts related to different stress-, hormone-, and reactive oxygen species (ROS)-response (left column) among putative electric- and calcium/reactive oxygen species-dependent transcripts significantly altered in their expression in plants during plant-to-plant communication. The number of transcripts is shown in parentheses. In support of data presented in Fig. 5.

**Dataset S11.** The different mutants used in this work.

**Dataset S12.** Description of transcripts in each node of the gene regulatory network analysis of the transcriptomic response of leaf S3 of receiver plants, following the application of excess light (EL) stress to the local leaf of transmitter plants, in the WT-WT-WT setup. In support of Fig. 8A.

**Dataset S13.** Description of transcripts in each node of the gene regulatory network analysis of the transcriptomic response of leaf S3 of receiver plants, following the application of excess light (EL) stress to the local leaf of transmitter plants, in the WT-*hpca1*-WT setup. In support of Fig. 8A.

**Dataset S14.** Description of transcripts in the gene regulatory network of IOS1 in the transcriptomic response of leaf S3 of receiver plants, following the application of excess light (EL) stress to the local leaf of transmitter plants, in the WT-WT-WT setup. In support of Fig. 8B.

**Dataset S15.** Description of transcripts in the gene regulatory network of BBE11 in the transcriptomic response of leaf S3 of receiver plants, following the application of excess light (EL) stress to the local leaf of transmitter plants, in the WT-WT-WT setup. In support of Fig. 8C.

